# Visualizing the Dynamics of DNA Replication and Repair at the Single-Molecule Molecule Level

**DOI:** 10.1101/2022.10.22.513350

**Authors:** Scott Berger, Gheorghe Chistol

## Abstract

During cell division, the genome of each eukaryotic cell is copied by thousands of replisomes – large protein complexes consisting of several dozen proteins. Recent studies suggest that the eukaryotic replisome is much more dynamic than previously thought. To directly visualize replisome dynamics in a physiological context, we recently developed a single-molecule approach for imaging replication proteins in *Xenopus* egg extracts. These extracts contain all the soluble nuclear proteins and faithfully recapitulate DNA replication and repair *in vitro*, serving as a powerful platform for studying the mechanisms of genome maintenance. Here we present detailed protocols for conducting single-molecule experiments in nuclear egg extracts and preparing key reagents. This workflow can be easily adapted to visualize the dynamics and function of other proteins implicated in DNA replication and repair.

## Background and Motivation

Eukaryotic genomes have dramatically increased in size during evolution, from 12 million base-pairs in yeast (haploid) to 3.2 billion base-pairs in humans (haploid) to a staggering 130-150 billion base-pairs in the largest-known genomes (Blommaert, 2020; Meyer et al., 2021). To efficiently copy their massive genomes, eukaryotes replicate DNA from thousands of sites called replication origins (~400 origins in yeast vs ~50000 origins in humans) (Ganier et al., 2019; Méchali, 2010). At least 50 different proteins act together to replicate DNA, forming a complex known as the <u>replisome</u> (Fig. 1) (Yao & O’Donnell, 2019; Yeeles et al., 2015). As a result, thousands of replisomes copy the genome in parallel during cell division. Fundamental aspects of the DNA replication mechanism remain poorly understood: How are replisomes assembled correctly during replication initiation? How do replisomes efficiently replicate DNA and robustly cope with roadblocks like DNA damage? How are replisomes dis-assembled upon completing replication? How is the activity of thousands of replisomes properly regulated to ensure the complete and timely replication of large eukaryotic genomes?

**Figure 1.**
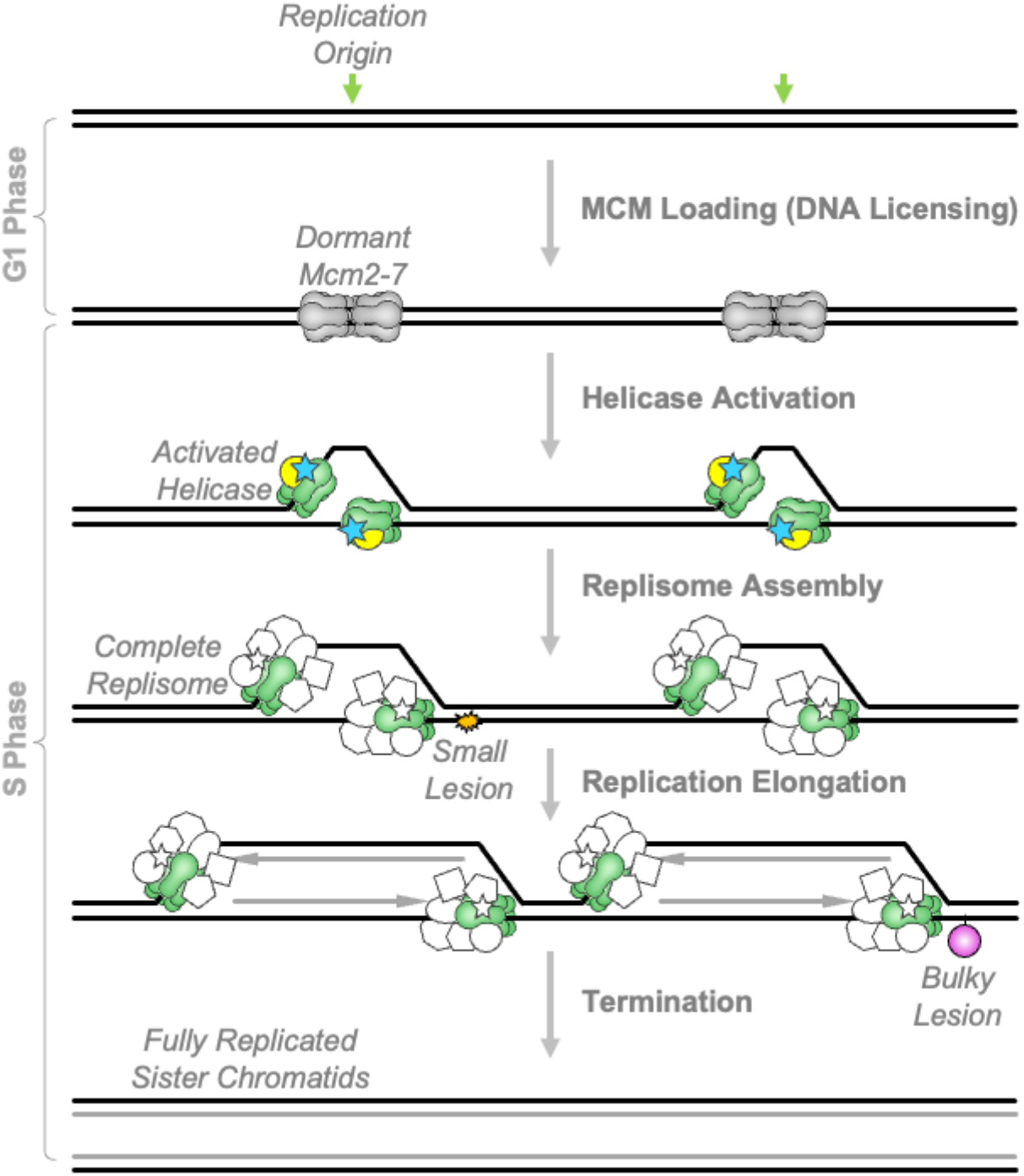
Overview of Eukaryotic DNA Replication. Licensing occurs in G1 phase. Two copies of the inactive replicative helicase Mcm2-7 are loaded onto double-stranded DNA at each replication origin in a head-to-head orientation. These double hexamers remain dormant until S phase, when they are activated resulting in the formation of mature replisomes and initiation of DNA replication. Upon completion of DNA replication, converging replisomes are unloaded. During elongation, replisomes encounter and cope with diverse DNA lesions – some of which are small enough to pass through the helicase, and others that are too big to do so.

DNA replication consists of four distinct phases (Fig. 1).

i. DNA licensing – wherein two inactive copies of the replicative helicase (Mcm2-7 complex) are loaded onto double stranded DNA at replication origins (also referred to “initiation zones” in metazoa) during G1 phase of the cell cycle (Bleichert, 2019; Frigola et al., 2013; Remus et al., 2009). Hence, these origins become “licensed” to replicate during the subsequent S phase (Blow & Laskey, 1988).
ii. Replication initiation – wherein a subset of replication origins is activated, or “fired”. This phase can be further subdivided into helicase activation and replisome assembly. During helicase activation, cell cycle-dependent kinases CDK and DDK promote the recruitment of two essential helicase subunits (GINS and Cdc45) to each Mcm2-7 complex, forming replicative helicase CMG (Cdc45+Mcm2-7+GINS) (De Jesús-Kim et al., 2021; Ilves et al., 2010; Muramatsu et al., 2010; Yeeles et al., 2015). CMG is a ring-shaped enzyme that unwinds DNA by threading one strand through its central pore (R. Georgescu et al., 2017; Li & O’Donnell, 2018). Numerous other replication proteins (polymerases, primase, processivity factors, structural proteins, ssDNA binding proteins) are then recruited to the helicase, forming a replisome. Notably, each origin that fires, gives rise to two replisomes that copy DNA bi-directionally from the origin (Coster & Diffley, 2017; Diffley, 2011).
iii. Replication elongation – wherein each replisome copies tens or hundreds of kilo-bases of DNA. During this phase the activity of the helicase is tightly coupled with that of the leading strand polymerase Pole (Byun et al., 2005).
iv. Replication termination – wherein replisomes from neighboring origins converge, complete replication, and are unloaded from chromatin in a manner that recycles replication proteins (Dewar et al., 2015; Dewar & Walter, 2017; Xia, 2021).

Over the past few decades, genetic studies in yeast outlined the core set of eukaryotic replication factors. Their homologs in higher eukaryotes were identified and validated in frogs, humans, mice, fruit flies, and nematodes. These efforts culminated in the biochemical reconstitution of yeast replication in yeast (Frigola et al., 2013; R. E. Georgescu et al., 2015; Remus et al., 2009; Yeeles et al., 2015), and recently – a partial reconstitution of human DNA replication (Baris et al., 2022). Advances in proteomic approaches revealed new components of the DNA replication and repair machineries (Alabert et al., 2014; Räschle et al., 2015; Sirbu et al., 2013). Finally, sequencing-based tools further advanced in our understanding of DNA replication, especially pertaining to the spatio-temporal regulation of replication origins (Claussin et al., 2022; Hennion et al., 2020; Macheret & Halazonetis, 2019; Marchal et al., 2018; Yu et al., 2014).

However, most current approaches sample thousands of replisomes per cell and millions of cells per sample. Although there exist strategies to synchronize cells or biochemical reactions, they still average the behavior of large numbers of replisomes in each sample, obscuring rare or asynchronous events.

Recent work suggests that replisomes are much more dynamic and plastic than previously thought (Lewis et al., 2017, 2020; Mueller et al., 2019). To study the molecular mechanisms that underlie DNA replication, we developed a novel single molecule approach whimsically named KEHRMIT (Kinetics of the Eukaryotic Helicase via Real-time Molecular Imaging and Tracking) (Sparks and Chistol et al., 2019).

We focused specifically on the replicative helicase for a few reasons. First, the helicase forms the core of the replisome around which all other proteins are assembled (Fig. 1) (Bai et al., 2017; Baretić et al., 2020). Second, helicase loading, activation, and unloading are strictly regulated to ensure that the genome is replicated correctly (O’Donnell et al., 2013), but this regulation remains poorly understood. Finally, helicase activity is critical for DNA damage signaling. Normally, the CMG helicase is physically and functionally coupled to the leading strand polymerase Polε. Many types of DNA damage “uncouple” these enzymes by inhibiting the polymerase but not the helicase (Byun et al., 2005). The uncoupled helicase continues to unwind double-stranded (dsDNA), converting it to single-stranded DNA (ssDNA) (Zeman & Cimprich, 2014). ssDNA activates the replication stress response, but is highly susceptible to breakage and threatens genomic integrity (Berti et al., 2020). How the uncoupled helicase is regulated to balance the benefits and risks of ssDNA remains unclear.

### Biomedical Relevance of Studying the Basic Mechanisms of DNA Replication and Repair

The replisome interfaces with hundreds of proteins involved in DNA metabolism, chromatin maintenance, and transcription (Dungrawala et al., 2015; Räschle et al., 2015). Studying replisome biology will advance our basic understanding of DNA replication and other cellular processes. There are numerous endogenous and exogenous sources of replication stress, including normal metabolism, DNA-bound protein roadblocks, transcription-replication conflicts, ultra-violet light, ionizing radiation, and chemotherapy drugs (Zeman & Cimprich, 2014). Therefore, understanding the replication stress response has key implications for human health (Berti et al., 2020). If cells fail to resolve replication stress, they experience a loss of genetic information, DNA breaks, chromosome mis-segregation, and other abnormalities known as genomic instability – a hallmark of cancer (Macheret & Halazonetis, 2015). Another hallmark of cancer is rapid cell proliferation enabled by oncogene overexpression, which drives fast DNA replication at the cost of elevated replication stress (Primo & Teixeira, 2020). Chemotherapy drugs, which remain the standard of care for many types of cancer, target tumor cells by causing excessive DNA damage or inhibiting DNA replication and repair (Kitao et al., 2018). However, these drugs also affect other cells that divide rapidly, causing severe side effects. Understanding the molecular underpinnings of DNA replication and replication stress response will enable the development of new diagnostic tools and therapies for cancer and other disorders (Berti et al., 2020).

### Overview of KEHRMIT

In KEHRMIT, DNA is stretched and tethered to a glass surface inside a flow cell (Fig. 1A), as previously described (Yardimci, Loveland, et al., 2012). This DNA substrate is then replicated in *Xenopus* egg extract immunodepleted of GINS (a helicase subunit) and supplemented with recombinant GINS labeled with a bright organic dye such as Alexa Fluor 647 (Fig. 1A, green stars). Thus, a fluorophore is incorporated into each helicase, acting as a tracking beacon. A Total Internal Reflection Fluorescence (TIRF) microscope is then used to simultaneously monitor the replication of several hundred DNA molecules with high spatial resolution (~150 base-pairs) and variable temporal resolution (0.1-100 sec). We recently used KEHRMIT to (1) dissect how the helicase copes with toxic DNA damage (Sparks and Chistol et al., 2019), (2) examine how the replisome is unloaded from chromatin during replication termination (Low et al., 2020), and (3) investigate the fate of the replisome when it encounters a nick or gap in the DNA template (K.B. Vrtis et al., 2021).

KEHRMIT experiments are performed in *Xenopus* egg extracts that contain the entire soluble proteome and efficiently recapitulate DNA replication and replication stress response (Walter et al., 1998). Due to their flexibility, *Xenopus* extracts have become a workhorse model system to study genome maintenance (Hoogenboom et al., 2017). For example, extracts can replicate genomic, linear, or circular DNA substrates. Moreover, any protein of interest can be immunodepleted, and the consequences of this “knock-out” can be investigated biochemically. Depleted extract can be “rescued” via addition of recombinant proteins. Hence, *Xenopus* extracts are an excellent system to study eukaryotic DNA replication proteins, many of which are essential and difficult to manipulate *in vivo*. This article describes in detail the KEHRMIT workflow summarized in Fig. 3.

### Other Single-Molecule Imaging Approaches Used in Conjunction w KERHMIT

#### (i) PhADE - PhotoActivation, Diffusion and Excitation

Although KEHRMIT enables the real-time imaging of the replicative helicase and its dynamics, it is often desirable to visualize the extent of replicated DNA. This provides independent validation that the KEHRMIT signal is indeed associated with bona fide DNA replication. To this end we visualize fluorescently labeled Fen1 (Flap Endonuclease 1), which binds to nascent lagging strands to facilitate Okazaki fragment maturation (Balakrishnan & Bambara, 2013). Specifically, we employ an approach called PhADE (PhotoActivation, Diffusion and Excitation), first proposed by Loveland et al., wherein Fen1 is fused to mKikGR (monomeric Kikume Green-Red) – a photoswitchable fluorescent protein that is constitutively green, but can be photoconverted into an orange-red fluorescent protein upon exposure to ultraviolet light (typically 405 nm) (Loveland et al., 2012).

Typically, Fen1^mKikGR^ is added to the single-molecule replication reaction to a final concentration of 500-1000 nM – roughly comparable to that of the endogenous protein (~300-500 nM as estimated previously in Wühr et al., 2014). These concentrations are much too high to enable conventional single-molecule detection via TIRF imaging (which requires fluorophore concentrations below ~50 nM). However, PhADE overcomes this limitation via a clever photoconversion-diffusion-excitation scheme illustrated in Fig. 2B and summarized in Table 1.

**Figure 2.**
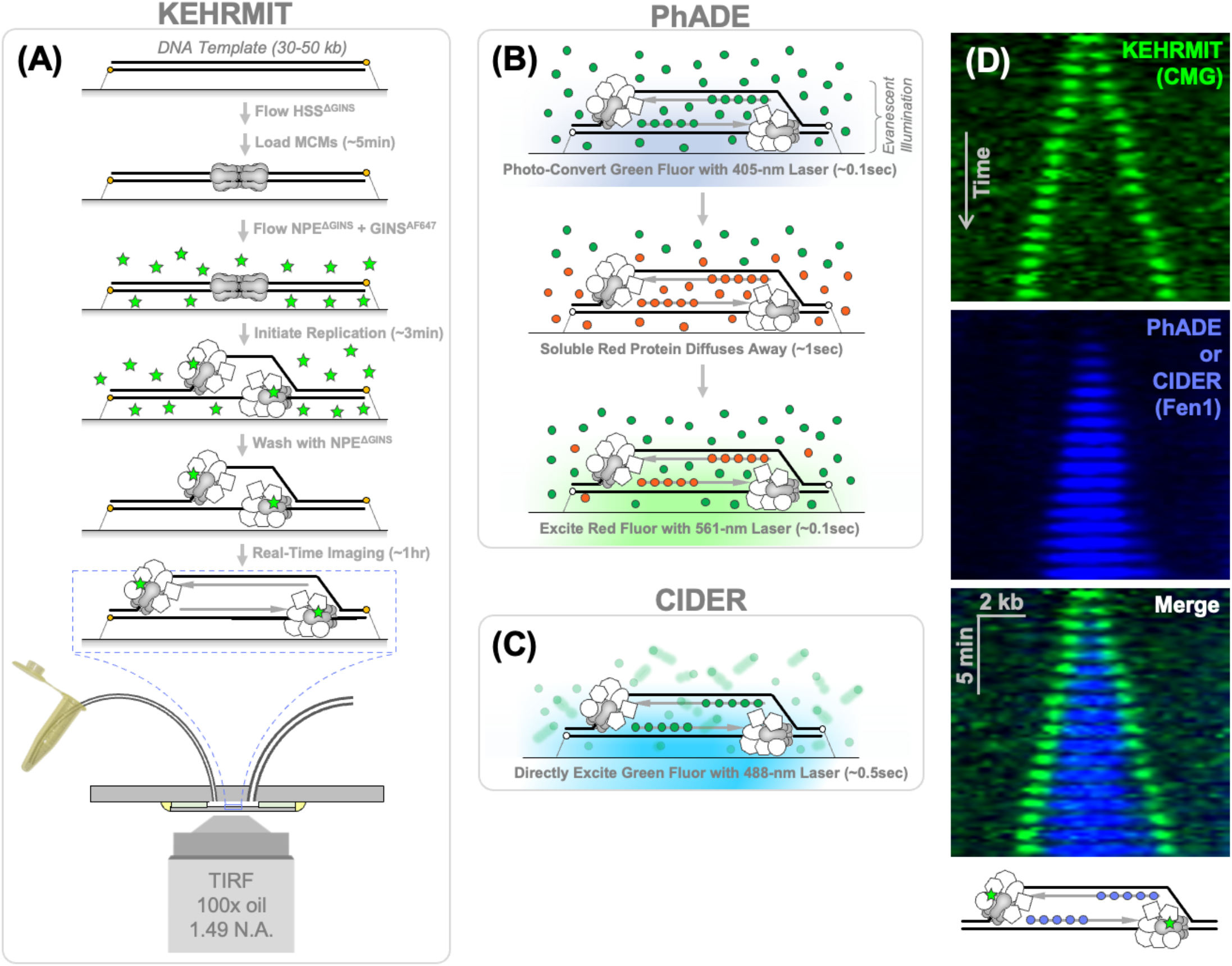
Overview of Single-Molecule Techniques to Study Replication in Xenopus Egg Extracts. In all three approaches (KEHRMIT, PhADE, and CIDER) a 30-50 kb DNA template is tethered to a coverslip in a microfluidic chamber. Next, HSS (extract that recapitulates the G1 phase) is drawn into the flow cell to license the DNA template. Next, NPE (extract that recapitulates S phase) is drawn in the flow cell to replicate DNA. **(A)** In KEHRMIT, NPE is immunodepleted of GINS and supplemented with recombinant GINS labeled with a fluorophore (AF647). The fluorescent GINS acts a beacon to directly monitor the spatio-temporal dynamics of individual replisomes. **(B-C)** In PhADE and CIDER, NPE is supplemented with Fen1 (or other proteins that associate with replication forks) fused to a fluorescent protein. In PhADE, a photo-convertible fluorescent protein (mKikGR) is used for rapid imaging of replication forks with high signal-to-noise ratio. In CIDER, photoactivation is skipped and the fluorescent protein is imaged directly via long exposures. **(D)** KEHRMIT and PhADE/CIDER may be used simultaneously for visualizing DNA replication dynamics.

**Figure 3.**
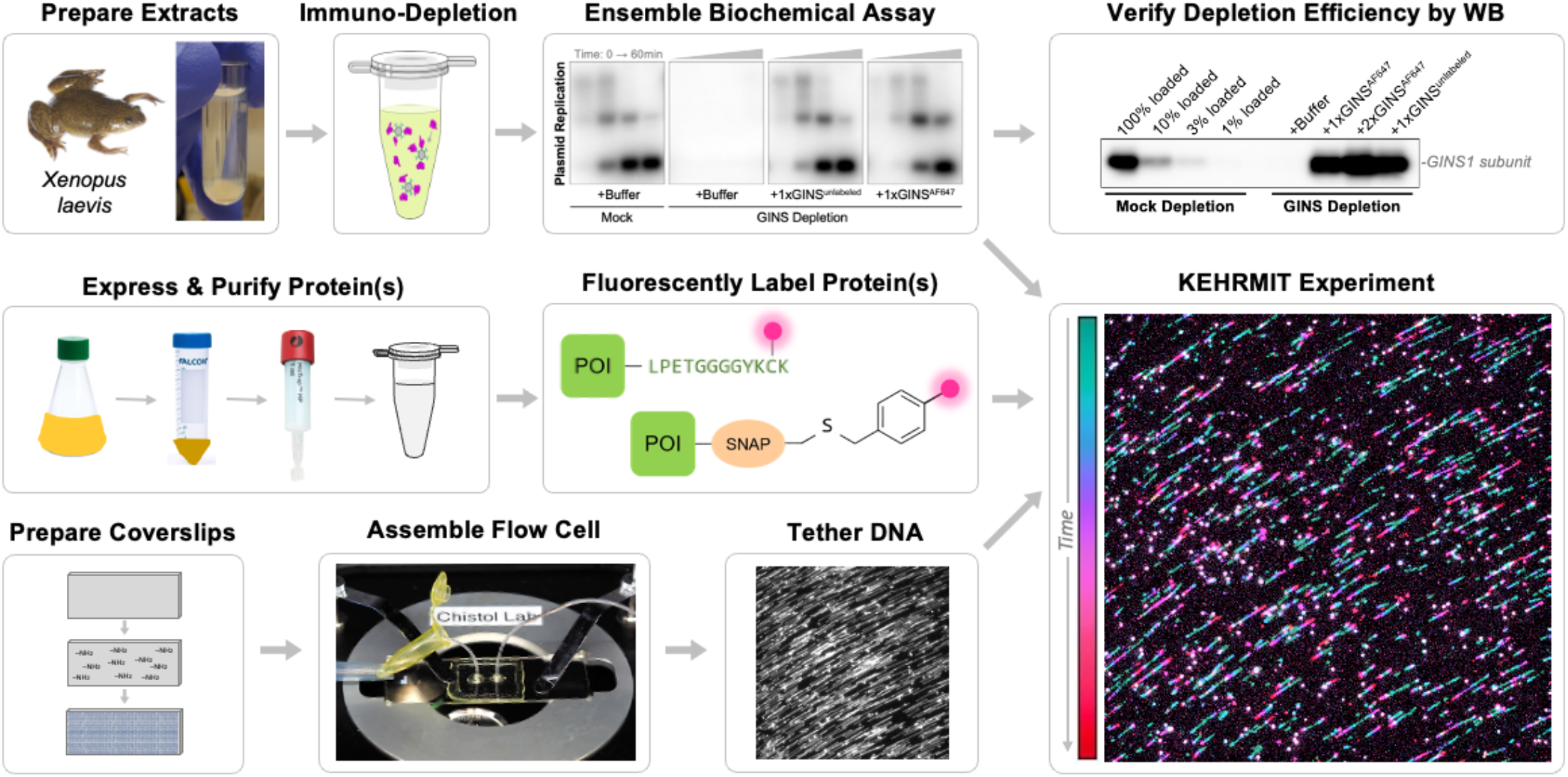
Overview of the KEHRMIT Workflow. Prior to conducting KEHRMIT experiments, extracts, antibodies, and recombinant proteins are extensively tested and validated via ensemble biochemical assays. In a KEHRMIT experiment, egg extracts are immunodepleted of one (or more) replisome proteins. The depleted extract is supplemented with recombinant fluorescently labeled protein. DNA substrates are tethered to a functionalized coverslip inside a microfluidic chamber. Egg extracts are flown into the microfluidic chamber to replicate DNA, and this process is monitored in a high throughput fashion using real time TIRF microscopy.

**Table 1:** Parameters for PhADE and _ CIDER Imaging Modalities.

| Mode | Imaging Phase | Laser Wavelength | Laser power out of 100x 1.4NA objective (through a square field aperture 100x100 microns) | Exposure Duration |
| --- | --- | --- | --- | --- |
| PhADE | Photoconversion of mKikG into mKikR within the evanescent field of TIRF illumination | 405 nm | 0.25 mW | 100 msec |
|  | Diffusion of soluble Fen1<sup>mKikR</sup> out of the evanescent field | none | - | >500 msec (usually 10-30 sec) |
|  | Excitation of Fen1<sup>mKikR</sup> bound to DNA substrates | 561 nm | 0.05 - 0.10 mW | 100 msec |
| CIDER | Direct excitation of Fen1<sup>mKikG</sup> (or Fen1<sup>mNeonGreen</sup>) | 488 nm | 0.05 - 0.10mM | 500 msec or longer |

First, Fen1^mKikG^ molecules within the evanescent field of TIRF illumination (~100 nm deep) are photoconverted from green from to the red form by a brief (typically 100ms) pulse of 405 nm light. Second, soluble Fen1^mKlkR^ molecules are allowed to diffuse away from the surface of the coverslip (typically 500ms or longer), while Fen1 ^mKlkR^ acting on nascent lagging strands remains bound to the tethered DNA substrate. Finally, Fen1^mKlkR^ molecules that decorate the replication bubble are visualized using a brief excitation pulse with the 561nm laser (~100ms). Although our laboratory employs PhADE only to measure the extent of replicated DNA, this strategy can be used to detect and count individual molecules (Loveland et al., 2012).

We have recently adapted PhADE to visualize RPA^mKikGR^. RPA is a single-stranded DNA binding protein that is recruited to ssDNA near Okazaki fragments during normal replication or to ssDNA generated during replication stress. In principle, PhADE could be used to visualize many other proteins involved in DNA replication which are present at high endogenous concentrations in the nucleus (PCNA, histones, etc).

#### (ii) CIDER: Continuous Imaging via Direct Excitation of Replication factors

One of the drawbacks of PhADE is that it “takes up” three standard spectral channels: the 405 nm channel required for photoconversion, the 488 nm channel where the signal is dominated by “naïve” mKikG that has not yet been photoconverted, and the 561 nm channel where the mKikR signal resides. Effectively this leaves 640 nm as the only spectral channel available for orthogonal single-molecule imaging. Although we successfully employed a combination of PhADE and KEHRMIT in two previous studies (Low et al., 2020; Sparks and Chistol et al., 2019), this strategy became limiting when it became imperative to visualize additional replication proteins (K.B. Vrtis et al., 2021).

In cases where chromatin is bound by many copies of a fluorescently labeled factor, it is possible to directly excite those fluorophores and visualize the DNA-bound proteins with high signal-to-noise even if the concentration of free labeled protein is very high (>1000 nM). This strategy was previously used to visualize Fen1 (K.B. Vrtis et al., 2021), RPA (Yardimci, Wang, et al., 2012), and ubiquitin (Lu et al., 2015) at physiological protein concentrations (~1000 nM) which are incompatible with conventional single-molecule TIRF imaging.

For convenience we internally refer to this imaging modality as CIDER – Continuous Imaging via Direct Excitation of Replication factors (Fig. 2C). For standard Fen1 or RPA experiments, CIDER provides a signal-to-noise performance comparable to that of PhADE while being simpler to perform and optimize imaging conditions. Our laboratory has recently entirely switched from PhADE to CIDER. Importantly, when using an EMCCD for CIDER imaging, low laser power is sufficient to achieve reasonable SNR because the primary source of noise is the diffusion of free fluorescent proteins. This noise is suppressed by simply averaging the signal for at least 500 msec (Table 1). Finally, note that PhADE may provide better SNR performance in cases where the number of DNA-bound proteins is very low.

### Examples of Discoveries Enabled by KEHRMIT

The transformative potential of KEHRMIT is illustrated by two studies summarized below (Low et al., 2020; Sparks and Chistol et al., 2019). Altogether, these experiments revealed that the helicase, and likely the rest of the replisome, are much more plastic than previously thought. Future single-molecule studies will be critical for understanding the causes and consequences of this plasticity. Importantly, each study contained a wealth of biochemical experiments (not discussed here) to verify and constrain our models, validate reagents, and corroborate the findings of single-molecule experiments. We are now dramatically expanding the capabilities of KEHRMIT by using multi-color imaging to simultaneously visualize the dynamics and function of numerous other replication proteins.

#### (i) Dissecting the Mechanism of DNA-Protein Crosslink Bypass and Repair

DNA-Protein cross links (DPCs) are toxic DNA lesions that form because of exposure to exogenous sources of damage (UV, ionizing radiation, chemotherapeutics) as well as endogenous sources (aldehydes resulting from normal metabolism or intermediates of failed DNA repair) (Weickert & Stingele, 2022). By forming a bulky roadblock on the DNA template DPCs represent a formidable challenge to both the transcription and replication machineries. In cycling cells, most DPCs are detected and repaired in a replication-dependent manner. Since many DPCs look indistinguishable from DNA-bound proteins they are only detected when a replisome collides with the lesion and cannot evict it from DNA (Duxin et al., 2014; Stingele et al., 2014).

Unlike many other DNA lesions, which are relatively small and are easily bypassed by the replicative helicase by threading DNA through the CMG central channel, DPCs are much too large to fit through CMG. It was previously thought that when the replisome encounters a DPC (Fig. 4A), the lesion stalls the helicase (Duxin et al., 2014; Stingele et al., 2014). It was then proposed that the stalled replication fork would trigger the proteolytic degradation of the lesion, and the helicase could subsequently resume unwinding DNA only after DPC proteolysis (Fig. 4A). This model predicts that when DPC proteolysis is blocked, it should stall the replication fork indefinitely.

**Figure 4.**
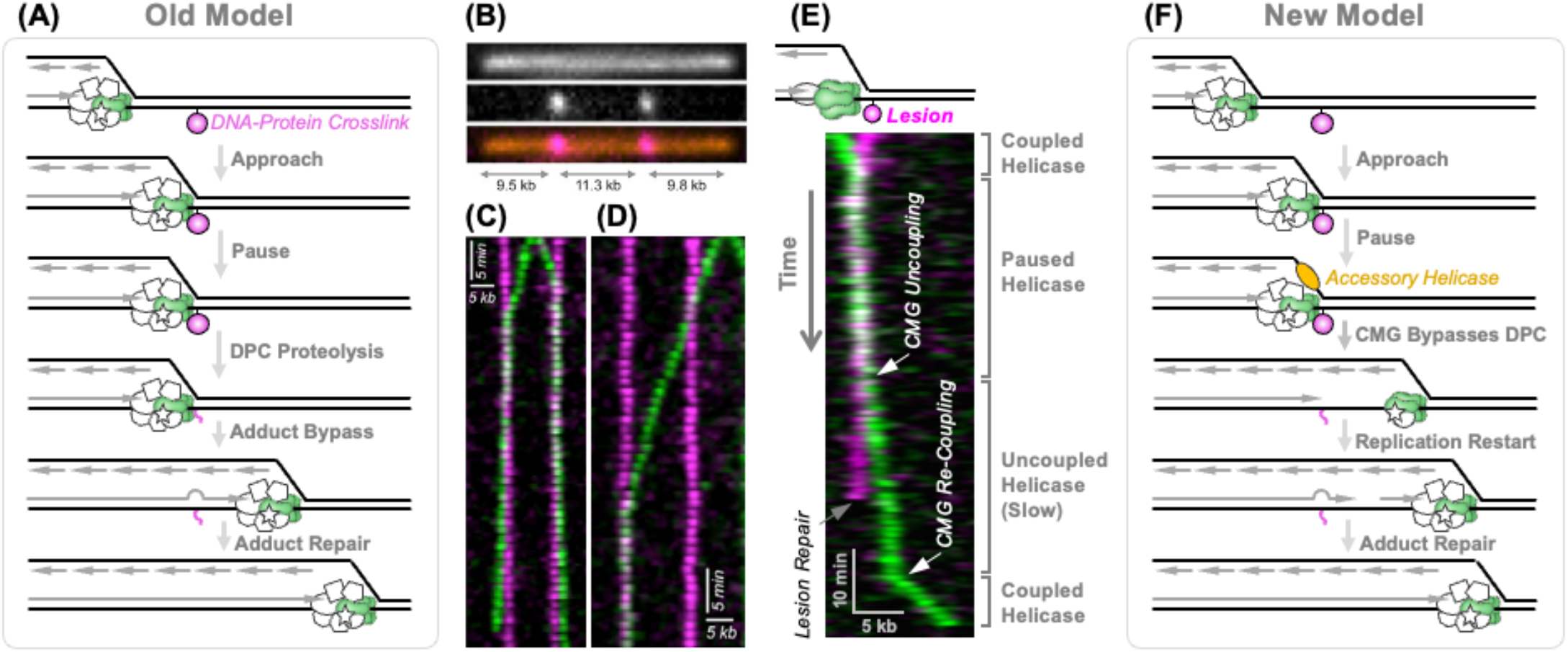
Using KEHRMIT to Dissect the Mechanism of DPC Repair. **(A)** Model of DPC repair prior to our single-molecule study. **(B)** Image of a model DNA substrate (red) containing two site-specific DPC lesions (magenta). **(C)** Kymogram of helicase (green) dynamics upon encountering DPCs (magenta) on the leading strand template. **(D)** Kymogram of helicase dynamics upon encountering DPCs on the lagging strand template. **(E)** Kymogram of helicase dynamics following an encounter with a DPC on the leading strand template which exhibits evidence for replication fork restart. **(F)** Model of DPC repair incorporating new insights gained from our KEHRMIT experiments.

To test this model, we engineered DPCs at specific locations on a DNA plasmid and used an ensemble biochemical assay to probe how far the newly synthesized leading DNA strand was extended. Normally, the leading strand is extended to within 30-40 nt from the DPC – corresponding to the footprint of the CMG helicase paused in front of the roadblock. After a brief delay, the nascent leading strand was extended all the way to the DPC indicating the CMG helicase “got out of the way” (presumably after the DPC was degraded) allowing the replicative polymerase to advance. However, when DPC proteolysis was prevented, the nascent leading strand advanced to the DPC location without delay, suggesting that the helicase was either “unloaded” from DNA, or that the helicase somehow “bypassed” the intact bulky lesion (Sparks and Chistol et al., 2019).

To unambiguously determine how the replicative helicase copes with a DPC we generated a custom linear DNA substrate with two DPCs engineered at specific locations in a specific orientation (Fig. 4B). The DPCs were created by cross linking the HpaII methyltransferase (45kDa) to unique sequences on the DNA. To visualize the DNA lesion, HpaII was fluorescently labeled with Alexa Fluor 568 (HpaII^AF568^) at the C-terminus using an engineered version of the bacterial enzyme Sortase A (Antos et al., 2017). KEHRMIT experiments were conducted using GINS^AF647^ to visualize the dynamics of the CMG helicase, and conditions were optimized to limit replication initiation to 1-2 origins per DNA molecule (which greatly simplified data analysis and interpretation) (Sparks and Chistol et al., 2019).

These experiments revealed several key insights. First, CMG could efficiently bypass DPCs under conditions when proteolysis was inhibited (Fig. 4C-D). Second, we observed CMG bypassing DPCs under “normal” conditions, with DPC destruction being observed after bypass (Fig. 4E). Third, we observed that it took ~25 min to degrade the DPC, but only ~15 min to bypass the DPC from the moment of CMG arrival, indicating that DPC bypass preceded DPC destruction (Sparks and Chistol et al., 2019). Fourth, the CMG helicase slowed down dramatically after bypassing an intact DPC on the leading strand template (Fig. 4C, E). This dramatic suppression of DNA unwinding enables the uncoupled helicase to generate sufficient ssDNA to activate the DNA damage checkpoint (Zeman & Cimprich, 2014) while minimizing the risk of breaking exposed ssDNA (Berti et al., 2020). We validated this idea by visualizing the dynamics of CMG in the presence of aphidicolin – an inhibitor of DNA synthesis that promotes helicase uncoupling (Sparks and Chistol et al., 2019). Fifth, uncoupled CMG resumed fast translocation sometime after the DPC lesion was degraded, suggesting that the CMG uncoupled from DNA synthesis and then re-coupled (Fig. 4E). Finally, we observed that CMG paused for ~15min at a DPC on the leading strand, and very briefly (~2min) at a DPC on the lagging strand (Fig. 4D), consistent with the fact that CMG translocates on the leading strand template, excluding the lagging strand template from its central pore (Fu et al., 2011a; Yuan et al., 2020).

In summary, KEHRMIT enabled us to directly visualize, for the first time, how the CMG helicase copes with a DNA lesion in a physiological setting. We demonstrated that CMG could bypass intact bulky lesions and that bypass serves as the trigger for lesion repair and degradation (Fig. 4F). It remains unclear how exactly CMG can bypass an intact DPC: CMG may be able to partially unfold the protein and thread it through its central pore, or the CMG ring may “crack” into a lock-washer configuration and thus slide past the DPC. These models remain to be tested explicitly, and single-molecule imaging will likely play an important role in that effort.

#### (ii) Dissecting the Mechanism of CMG Unloading During Replication Termination

Upon completion of DNA replication, the replisomes and associated proteins are unloaded in a highly regulated manner (Dewar & Walter, 2017; Xia, 2021). Until recently, replication termination remained poorly understood due to the asynchronous timing and distributed spacing of replication termination events. Advances in biochemical reconstitution of yeast replication, and the development of new tools for synchronizing replication termination enabled the systematic dissection of this previously understudied stage of DNA replication (Deegan et al., 2019; Dewar et al., 2015; Heintzman et al., 2019; Maric et al., 2014). It was discovered that during replication termination, a E3 ubiquitin ligase (Scf^Dia2^ in yeast and CUL2^LRR1^ in vertebrates) poly-ubiquitylates the Mcm7 subunit of the CMG helicase, which is subsequently extracted from chromatin by p97/VCP (Fig. 5A). The remaining replisome subunits dissociate from chromatin following helicase unloading.

**Figure 5.**
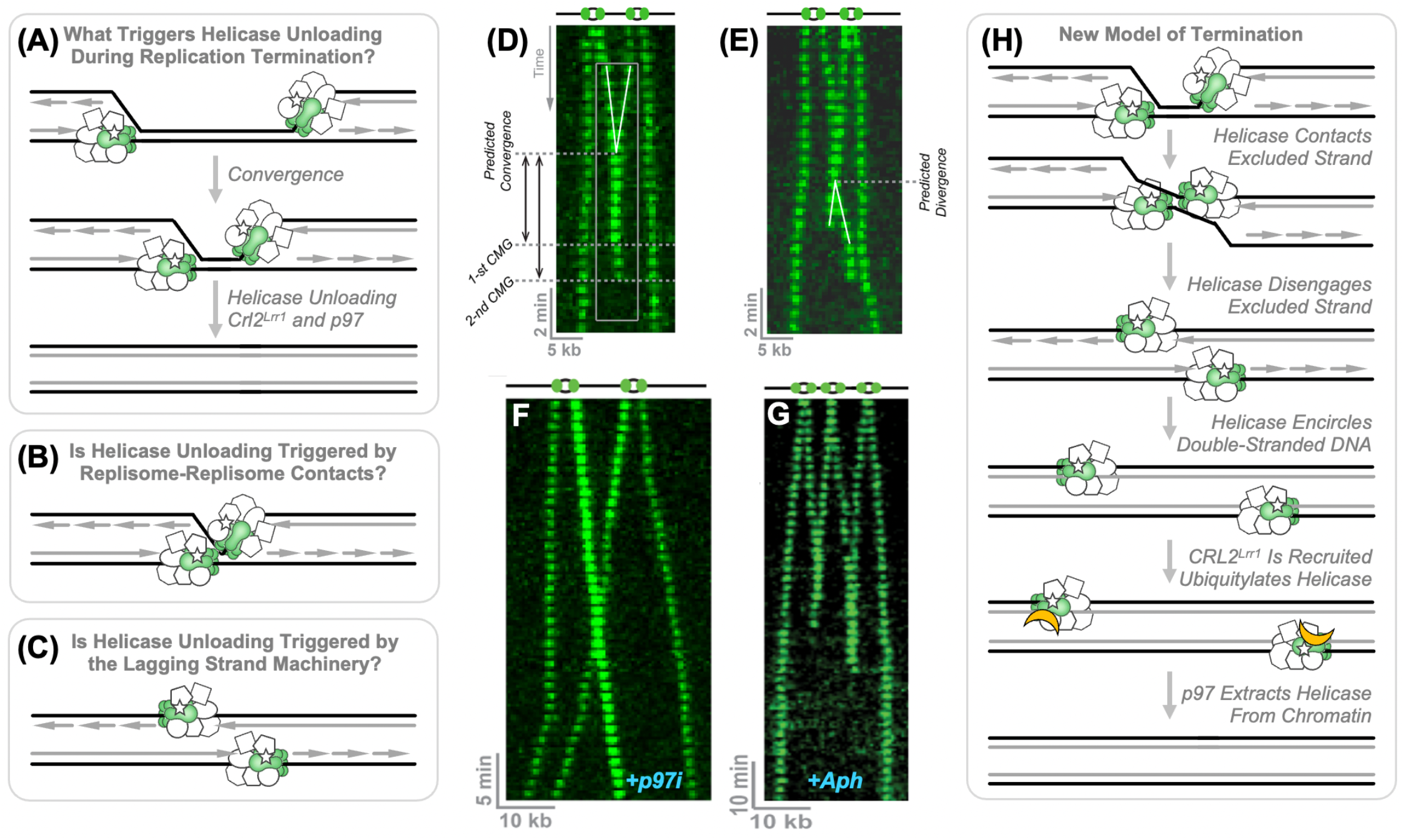
Using KEHRMIT to Dissect the Mechanism of Replication Termination. **(A)** Summary of our understanding of termination prior to our single-molecule studies. **(B-C)** Potential models to explain how replisomes are specifically unloaded during replication termination. **(D-E)** Example KEHRMIT kymograms visualizing, for the first time, helicase unloading after replisome convergence. **(F)** Example kymogram illustrating that helicases are no longer unloaded after replisome convergence in the presence of a p97 inhibitor. **(G)** Example kymogram illustrating that helicases are still unloaded in the absence of DNA replication (achieved by inhibiting DNA polymerases with Aphidicolin). **(H)** Model of replication termination incorporating new insights gained from our KEHRMIT experiments.

However, one key fundamental question eluded all previous genetic, genomic, proteomic, cellular, and biochemical studies - what is the molecular signal that triggers helicase unloading during termination (Fig. 5B-C)? Importantly, this signal must be highly specific to replication termination as premature helicase unloading would lead to DNA under-replication, and delayed unloading could cause DNA re-replication or may result in replication proteins remaining on DNA and acting as roadblocks to other chromatin-dependent processes.

To understand the exact nature of the molecular trigger for CMG unloading, we recapitulated the process on tethered lambda DNA (48.5kb) in a flow cell in *Xenopus* egg extracts and monitored helicase dynamics and unloading using our standard KEHRMIT assay (Fig. 2A). To maximize the likelihood of observing replication termination events, we modified the assay to yield ~2-4 origin firing events per DNA template: (i) we increased the licensing duration in HSS and (ii) we increased the replication initiation duration in NPE prior to washing off excess GINS^AF647^ (Low et al., 2020). We readily observed replication termination events which resulted in rapid (1-3) min unloading of the two CMG helicases after the convergence of two replisomes (Fig. 5D). Importantly, we observed that in a significant portion (~30%) of termination events, the two converged CMGs passed each other and were unloaded shortly after that (Fig. 5E). In either case, the two CMG molecules were unloaded at different times – i.e. unloading of the two replisomes is independent of each other and is not concerted (Fig. 5D-E).

We repeated the experiment in the presence of a p97 inhibitor (p97i) that prevented CMG unloading and observed that after converging and passing each other, helicases are not unloaded, but are able to travel long distances on DNA (Fig. 5F). Differences in the fluorophore brightness enabled us to unambiguously determine that converging CMGs did indeed pass each other (Fig. 5F). We repeated the experiments with Fen1^mKikGR^ (as a marker of lagging strand synthesis) and verified that diverging CMGs are not accompanied by DNA rereplication (Low et al., 2020). This led us to conclude that after replisomes converge, the helicases do not continue unwinding newly synthesized DNA, but rather continue translocating onto dsDNA which can fit though the central pore of CMG (a model proposed earlier by Dewar et al., 2015). Importantly, such a dsDNA translocation mode has been previously observed with recombinant CMGs (Kaplan et al., 2003; Wasserman et al., 2019).

At this point, it was tempting to hypothesize that CMG encircling dsDNA represents the molecular signal that triggers CMG unloading during termination. To test this model, we performed the KEHRMIT termination experiment in conditions that strongly inhibit DNA synthesis – i.e. in the presence of high concentrations of aphidicolin. If the model proposed above is true, then aphidicolin should strongly inhibit the unloading of converged CMGs. However, KEHRMIT experiments clearly showed that uncoupled CMGs are robustly unloaded from chromatin in the absence of DNA synthesis (Fig. 5G).

After ruling out several previously proposed models, we considered a new model where the trigger for CMG unloading during replication termination is the loss of interaction between the excluded DNA strand (lagging strand template) and the outer surface of CMG (Fig. 5H). In this simplified model, the E3 ligase needs to dock to the outer surface of CMG or needs access to these residues to ubiquitylate CMG. However, during normal replication this outer surface of CMG is “blocked” by the excluded DNA strand. Upon replisome convergence during termination, this interaction is abolished, and the E3 ligase can now access CMG and ubiquitylate it. This hypothesis is supported by another study enabled by KEHRMIT (Kyle B Vrtis et al., 2021). Gratifyingly, the model proposed by us was later corroborated by high resolution structures of CMG with CUL2^LRR1^ (Jenkyn-Bedford et al., 2021). Importantly, this beautiful structural study shows that the termination mechanism is conserved between yeast and humans. It is likely that other replisome subunits play additional roles in recruiting the ligase and regulating its function. This is an active area of inquiry and will likely involve a combination of biochemical reconstitution, structural studies, and single-molecule approaches.

## METHODS

### Extract Preparation

In KEHRMIT, PhADE, and CIDER experiments, two distinct *Xenopus* egg extracts are used. Both extracts are prepared by crushing fresh eggs via centrifugation and harvesting the crude cytoplasmic fraction (Fig. 6). The first extract, called High Speed Supernatant (HSS), recapitulates the G1 phase of the cell cycle and can be used to license circular DNA, linear DNA, or sperm chromatin (Lebofsky et al., 2009; Sparks & Walter, 2019; Walter et al., 1998). The second extract, called Nucleo-Plasmic Extract (NPE), is prepared by adding purified sperm chromatin to crude cytoplasmic extract, inducing the formation of nuclei. These nuclei are harvested via centrifugation and fractionated via ultra-centrifugation to obtain DNA-free and membrane-free NPE. This extract recapitulates the S phase of the cell cycle and contains a high concentration of replication proteins. NPE is used to initiate the replication of licensed DNA, supports efficient DNA replication elongation and replication termination as well as DNA repair and chromatin maintenance (Hoogenboom et al., 2017). We prepare sperm chromatin, HSS, and NPE following detailed protocols presented in Lebofsky et. al. 2009 and J. L. Sparks et al., 2019 with a few changes summarized below.

**Figure 6.**
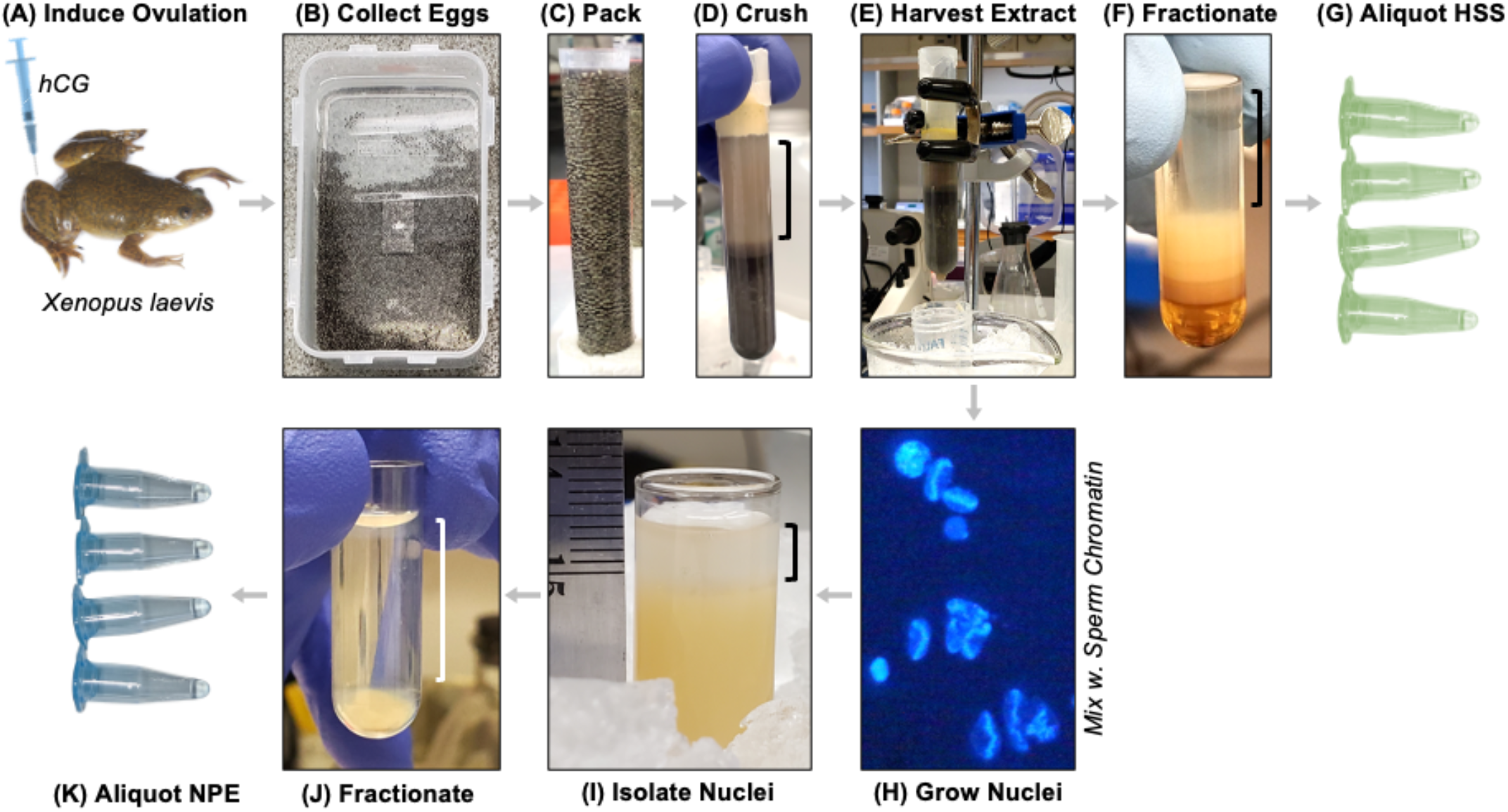
Overview of the Workflow for Preparing Xenopus Egg Extracts HSS and NPE. Eggs are harvested from *Xenopus laevis* females **(A-B)**, packed into culture tubes **(C)**, crushed via centrifugation **(D)**, crude cytoplasmic extract is harvested **(E)** and fractionated **(F)** to yield HSS **(G)** – a cytoplasmic extract that recapitulates the G1 phase. Sperm chromatin is mixed with crude cytoplasmic extract to form nuclei **(H)**, which are then isolated via centrifugation **(I)**, and fractionated **(J)** to yield NPE – a nucleoplasmic extract that recapitulates the S phase **(K)**.

#### Inducing Ovulation

Lyophilized recombinant human Chorionic Gonadotropin (hCG) (Chorulon Merck #140-927) is reconstituted with the included sterile 1x PBS. Injections are performed using a 1 mL slip tip syringe with a 27Gx1/2 needle (Becton Dickinson #305109). 5-7 days prior to extract preparation, each frog is primed with 75 IU (international units) of hCG (200 μL of 375 IU/mL). One day before extract preparation ovulation is induced by injecting each frog with 660 IUs of hCG (200 μL of 3300 IU/mL). Injected frogs are isolated overnight in individual plastic containers with 2L of 100 mM NaCl (the salt prevents the eggs from sticking to the walls of the container) (Fig 7A). This enables the researcher to inspect each clutch of eggs separately. Any leftover hCG solution is flash-frozen in liquid nitrogen and stored @ −80 °C for up to a year.

**Figure 7.**
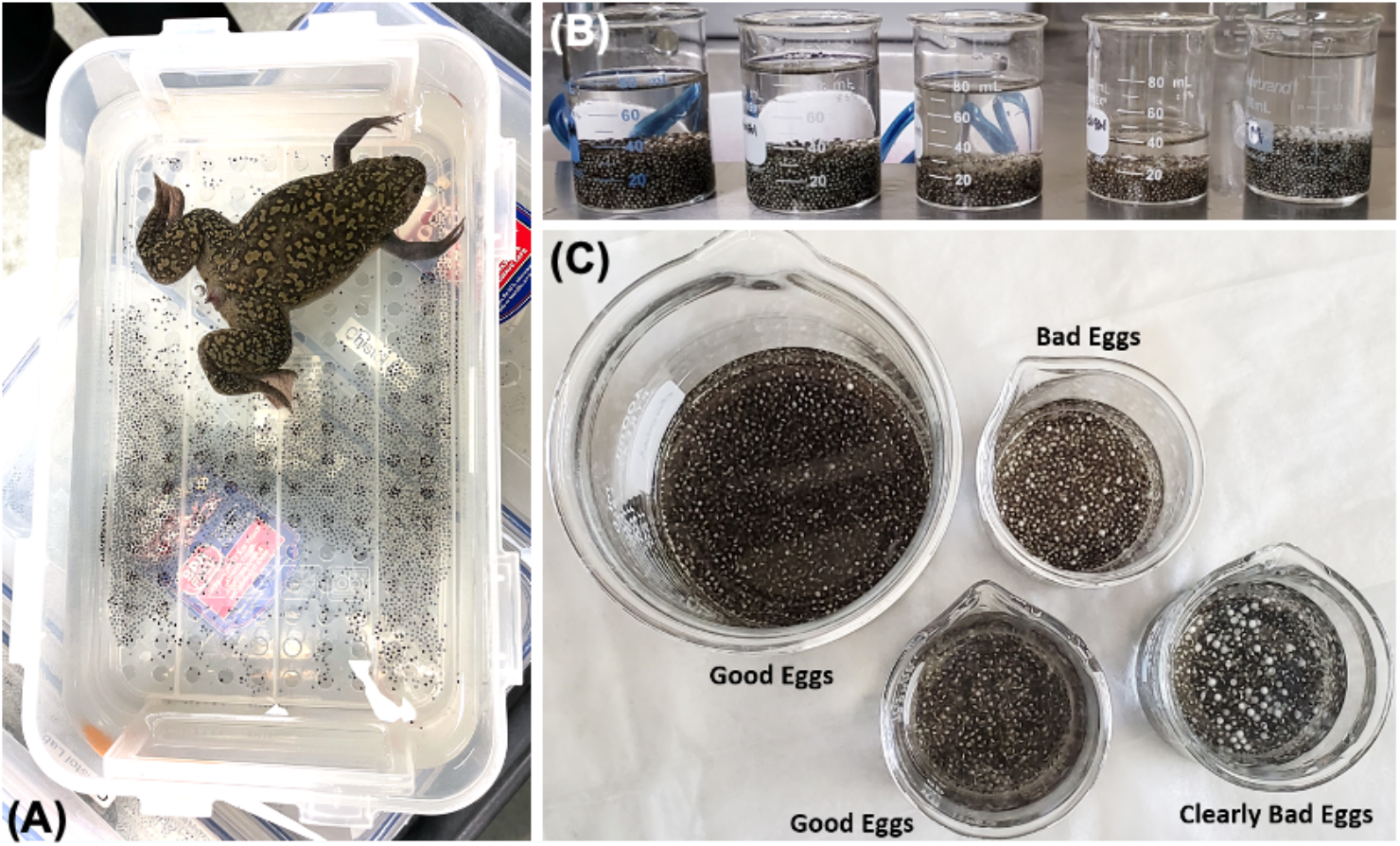
Guidelines for Selecting Oocytes for Egg Extract Preparation. **(A)** Frogs lay eggs overnight in individual containers with holes drilled in the lid for air exchange. We use a plastic container (Lock & Lock Airtight 121 oz) with a bottom drain insert that separates the eggs from the frog (frogs often eat their own eggs). A good clutch has clear water, eggs that do not stick to each other (or the container), has no white apoptotic eggs (or perhaps has 1-2 such eggs that can be picked out with a transfer pipette). **(B)** Eggs from each frog are collected into small beakers for inspection and yield estimates. **(C)** Overhead view of beakers with two good egg clutches, and two bad egg clutches (more than a few white apoptotic eggs).

#### Harvesting Eggs

Eggs are harvested after 20-21 hrs following the hCG injection. The quality and freshness of eggs is critical for optimal extract activity. Eggs harvested from each frog are collected in 100mL glass beakers and assessed separately (Fig. 7B). Typical yields range from 20mL to 40ml of eggs per frog. Frogs that lay less than 10 mL of eggs are euthanized. The following egg clutches are discarded and corresponding frogs are euthanized: (i) clutches with more than a few apoptotic white eggs (Fig. 7C), (ii) clutches with stringy eggs, (iii) clutches with very cloudy water (typically caused by egg lysis).

#### Egg Packing & Crushing

For both HSS and NPE preparations, eggs are packed at 100xg for 1 min at room temperature (Fig. 6C) in an ELMI CM-7S centrifuge with the 6M.02 swing bucket rotor. Eggs are crushed (Fig. 6D) at 15,000 xg for 20 min in a Beckman Avanti JE centrifuge with a JS13.1 swinging-bucket rotor in round-bottom 17×100 mm culture tubes (Corning #352059) held in custom 3D-printed nylon adapters (Shapeways). Prior to starting preparation, the rotor and adapters should be pre-equilibrated to room temperature and the centrifuge should be pre-cooled to 4C. The rotor is installed into the pre-cooled centrifuge immediately before starting the centrifugation, allowing the eggs to cool down gradually as they are crushed.

#### HSS Preparation

Eggs from 5-6 frogs are harvested 20 hours after hCG injection. Fractionation via ultracentrifugation (Fig. 6F) is carried out in a Beckman Optima MAX-XP tabletop ultracentrifuge in a TLS-55 swinging-bucket rotor at 4C at 55,000 rpm for 2 hrs. Extract is aliquoted into 0.2 mL PCR tubes as 45 μL aliquots (Fig. 6G), snap-frozen in liquid nitrogen, and stored at −80C for up to one year. For optimal extract performance, the entire protocol from the egg de-jellying to snap freezing should take no more than 4 hrs.

#### Sperm Chromatin Preparation

Testes are surgically removed from 10-14 euthanized male frogs. Frogs are euthanized by immersion for 10 min in 1 L water + 5 g Tricaine-S (Pentair via Fisher Scientific #NC0342409) + 10 g sodium bicarbonate (Fisher Scientific # MK-7396-500), followed by cervical dislocation.

#### NPE Preparation

Eggs from 10-15 are harvested 20-21 hours after the injection with hCG. Nuclei are isolated via centrifugation (Fig. 6I) at 15,000xg for 3 min at 4C in a Beckman Avanti JE centrifuge with a JS13.1 swinging-bucket rotor. Nucleoplasmic extract is fractionated via ultracentrifugation (Fig. 6J) and is then aliquoted into 0.2 mL PCR tubes as 11 μL aliquots (Fig. 6K), snap-frozen in liquid nitrogen, and stored at −80C for up to one year.

### Preparing Biotinylated DNA Substrates

Previous single-molecule studies of DNA replication with *Xenopus* egg extracts or purified proteins used a combination of single-tethered or double-tethered DNA (Duzdevich et al., 2015; Fu et al., 2011b; Gruszka et al., 2020; Lewis et al., 2017, 2020; Wasserman et al., 2019; Yardimci et al., 2010; Yardimci, Loveland, et al., 2012). Because single-tethered DNA molecules are rapidly chromatinized in egg extract and become highly condensed, they are not suitable for spatially resolved real-time imaging. Therefore, we exclusively use double-tethered DNA molecules 20-50 kb long. These DNA substrates are stretched using laminar flow and anchored to the glass coverslip via streptavidin-biotin linkages which maintain the DNA in an extended conformation for the duration of the experiment (typically stretched to 70 −90% of its contour length). Depending on the amount of slack in the double-tethered DNAs, a few to several nucleosomes may be loaded in extract until the DNA becomes taut (Gruszka et al., 2020).

Fig. 8 outlines several strategies for preparing DNA substrates for single molecule imaging. Below we present a detailed protocol for preparing long linear DNA substrates (Fig. 8A) as used in our recent studies (Low et al., 2020; Sparks and Chistol et al., 2019). The protocol for preparing substrates with site-specific DNA-Protein Crosslinks (DPCs) is presented in J. L. Sparks et al., 2019 (Fig. 8B-C). The workflow for preparing substrates with modified DNA inserts (Fig. 8E) is presented in Liu et al., 2015. Additional strategies for preparing custom DNA substrates can be found in Belan et al., 2021; Kim et al., 2017; Mueller et al., 2020.

**Figure 8.**
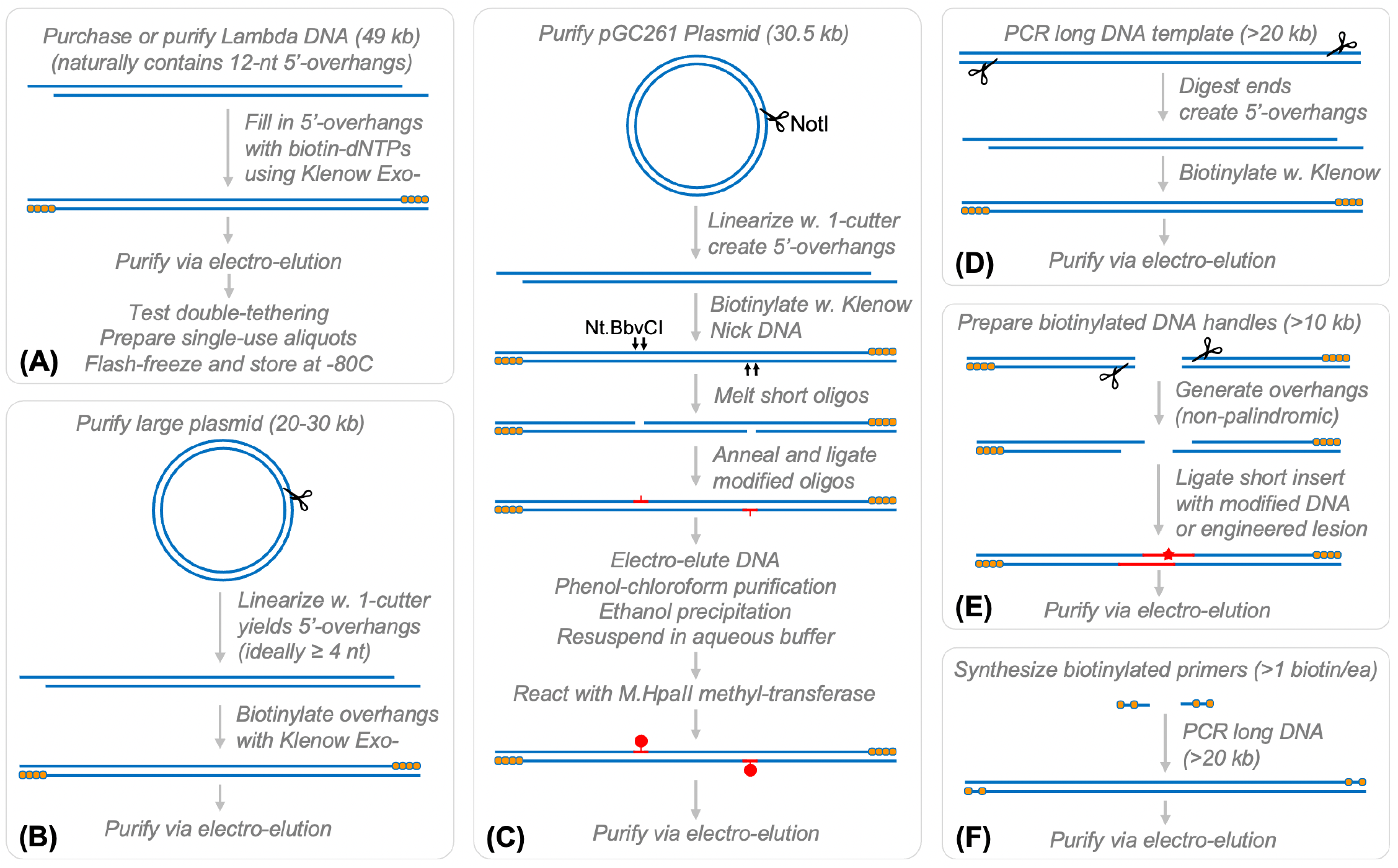
Diverse Strategies for Preparing DNA Templates for Single-Molecule Experiments. **(A)** Lambda phage DNA overhangs are filled-in with biotinylated dNTPs using the Klenow polymerase. **(B)** A large plasmid is linearized with a single restriction endonuclease, and the 5’ overhangs are biotinylated with Klenow. **(C)** Workflow for generating DNA templates with site-specific DPCs. **(D)** Another method to generate long DNA templates is to use a long-range DNA polymerase to PCR large (>20 kb) DNA fragments, digest their ends, and fill-in the 5’ overhangs as in panels (A-B). **(E)** A common strategy to introduce a site-specific lesion is to PCR long (>10 kb) DNA “handles” that can be ligated onto a small double-stranded DNA oligo with the engineered lesion. For best results this approach requires non-palindromic restriction enzymes. **(F)** Long DNA templated with modified ends can also be generated by performing long-range PCR with using primers that have been chemically modified (for example biotinylated).

### Double-Biotinylation of Lambda DNA

Lambda DNA is a convenient 48.5kb DNA substrate (Fig. 8A) that can be purified in house or purchased from a commercial supplier (NEB #N3011S). Lambda DNA contains 12-nt 5’ single-stranded DNA overhangs at each end (so called “cos-sites” 5’-GGGCGGCGACCT-3’ and 5’-AGGTCGCCGCCC-3’). Cos sites can be functionalized by ligating complementary 12-nt biotinylated oligos (Yardimci, Loveland, et al., 2012). We biotinylate lambda DNA by filling in the cos sites with biotin-dNTPs using the Klenow Exo-polymerase fragment. This approach is generalizable to other long DNA substrates for which 5’-overhangs can be generated (Fig. 8B). For example, we sometimes linearize 20-40kb plasmids with a single-cutter restriction enzyme that produces 4-nt 5’ overhangs that are subsequently filled in.

Following the Klenow fill-in, functionalized DNA is separated via electrophoresis and electro-eluted from the gel slice. This step is necessary to remove the vast excess of biotinylated nucleotides that would otherwise block all streptavidin binding sites on the surface of the microfluidic flow cell. By mixing and matching regular dNTPs with biotinylated dNTPs, we control the number of biotinylated nucleotides at each end of the DNA substrate. For example, in the case of lambda DNA one could carry out the reaction with biotin-dGTP + dTTP + dATP + dCTP to insert several biotins at each terminus, or biotin-dATP + dTTP + dGTP + dCTP to generate a single biotin at each DNA end.

1. Fill in 5’ single-stranded DNA overhang with dNTPs and biotin-dNTPs
  ○ 40 μL Lambda DNA @ 500 ng/μL (NEB #N3011S)
  ○ 6 μL 10x NEB2 Buffer
  ○ 2 μL 1mM dATP (Fisher #R0141) or biotin-dATP (Fisher #19524016) (33 μM f.c.)
  ○ 2 μL 1mM dCTP (Fisher #R0151) or biotin-dCTP (Axxora #JBS-NU-809-BIO16) (33 μM f.c.)
  ○ 2 μL 1mM dGTP (Fisher #R0161) or biotin-dGTP (Perkin Elmer #NEL541001EA) (33 μM f.c.)
  ○ 2 μL 1mM dTTP (Fisher #R0171) or biotin-dUTP (Fisher #R0081) (33 μM f.c.)
  ○ 2 μL ultrapure water
  ○ 4 μL Klenow Exo-enzyme (NEB #M0212S) μL *Incubate at 37°C for 30 min*,
  ○ Add 2 μL 0.5M EDTA to stop reaction
2. Run reaction on 0.7% Agarose gel (Fisher Scientific #03-500-523) pre-stained with SYBR Safe (Invitrogen #S33102) in 1xTAE buffer (Boston BioProducts #BM-250) at 5V/cm for 90 min. Excise the DNA band from the gel on a blue trans-illuminator and electroelute DNA in 1xTAE buffer at 5V/cm for 60 min in 6-8 kD cutoff dialysis tubing (Spectrum #132650).
3. Carefully collect eluate, avoid pipet-mixing to prevent shearing long DNA, estimate DNA concentration using a NanoDrop or via electrophoresis. Typically, we recover 50-80% of all DNA.
4. Test DNA for correct tethering in a single-molecule flow cell. Determine the correct amount of DNA for optimal density of double-tethered DNA.
5. Prepare single-use aliquots in 0.2 mL PCR tubes (usually 50-100 ng/ea) and flash-freeze in liquid nitrogen. Store at −80C for up to several years.

### Preparing PEG-Functionalized Coverslips

Coverslips functionalized with PEG and biotin-PEG are prepared following a previously published protocol with a few modifications (Tanner et al., 2009). The workflow is outlined in Fig. 9.

**Figure 9.**
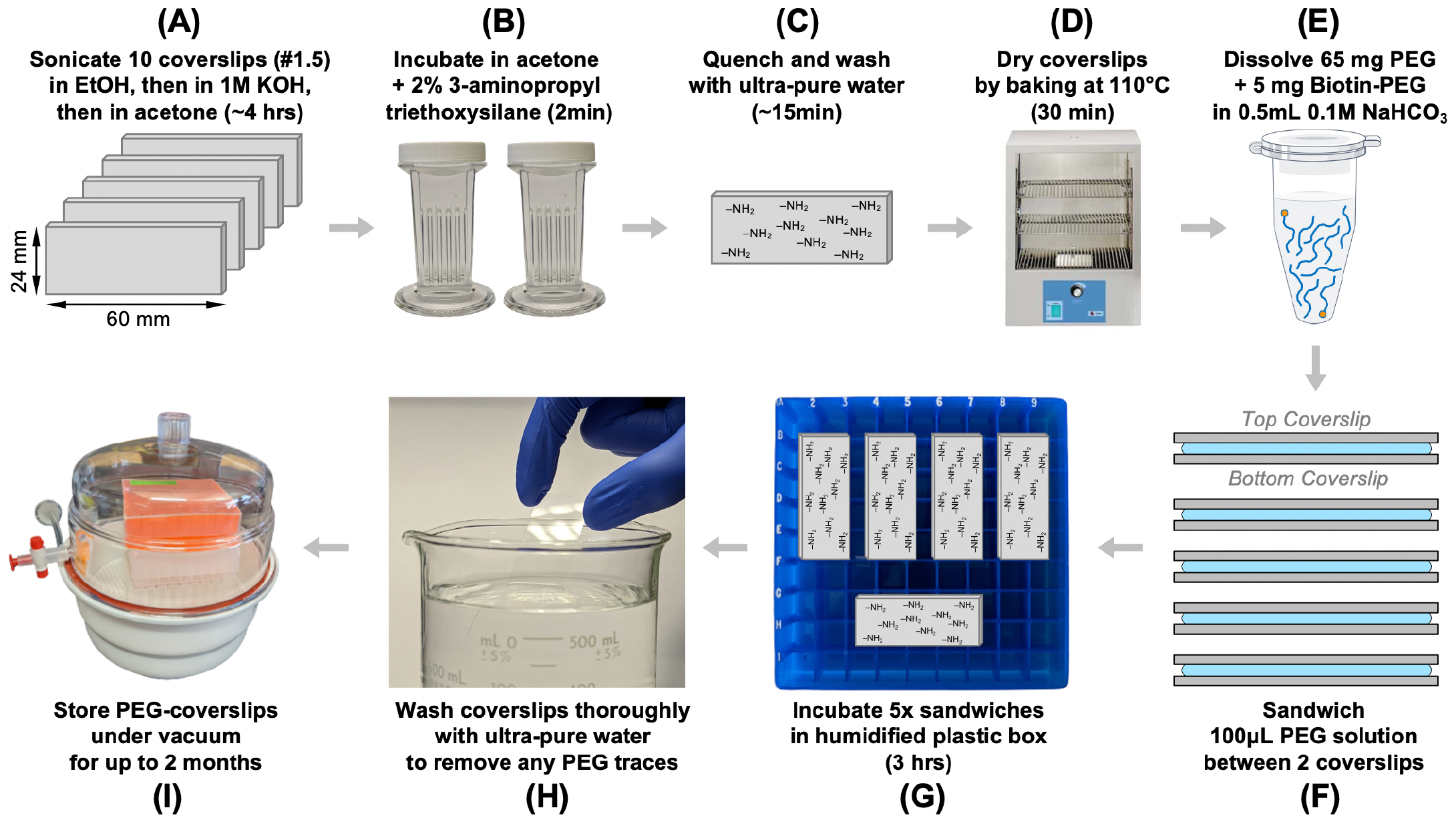
Preparing Functionalized Coverslips for KEHRMIT Experiments. **(A)** 24 x 60 mm coverslips are sonicated three times in EtOH and 1 M KOH. **(B)** Coverslips are then incubated in 2% 3-aminopropyl-triethoxysilane for 2 min while gently mixing. **(C)** The reaction is quenched by adding ultra-pure water for ~15 min. **(D)** Coverslips are then dried by baking at 100 °C for 30 min. **(E)** A PEG + biotin-PEG solution is prepared. **(F-G)** Coverslips are incubated with the PEG solution by assembling “sandwiches”. **(H)** Coverslips are then extensively washed with ultrapure water and stored under vacuum for up to 2 months **(I)**.

#### Materials

- 24 mm x 60 mm #1.5 coverslips (VWR #48393-106)
- Glass staining jars (DWK Life Sciences #900570)
- 5-mL glass serological pipette (plastic pipettes will react with silane)
- Silane (3-Aminopropyltriethoxysilane) (Sigma #A3648-100mL), store at 4C in inert gas (typically argon), in a container with desiccant, purchase fresh every 12 months
- Biotin-PEG-SVA MW5000 (Laysan Bio #Biotin-PEG-SVA-5000-100mg), store at −20C in argon, inside a container with desiccant, purchase fresh every 12 months
- mPEG-SVA MW5000 (Laysan Bio #MPEG-SVA-5000-1g), store at −20C in argon, inside a container with desiccant, purchase fresh every 12 months
- Acetone (Fisher Scientific #A18P-4)
- 1M KOH (0.45μm filtered)
- 200 proof ethanol (Gold Shield #412811)
- Oven (Fisher Scientific #11-475-152)
- Thermocouple for monitoring oven temperature (Amazon #B071V7T6TZ)
- Stainless steel uncoated chef’s cooling rack 8×11 inches (Amazon #B06Y5F3NGY)
- Ultrasonic water bath (VWR #97043-968)
- Vacuum desiccator dome (Ted Pella #2246)
- Argon gas

#### Clean Coverslips

1. Place 5 coverslips per staining jar, 10 coverslips total in 2 jars
2. Fill jars with ethanol, sonicate in the water bath for 30 minutes,
3. Pour off ethanol, rinse with ultrapure MilliQ water
4. Fill jars with 1M KOH solution, sonicate for 30 minutes
5. Pour off KOH, rinse with ultrapure MilliQ water
6. Repeat steps 2-5 for a total of three ethanol-KOH cleaning cycles
7. *Coverslips can be stored overnight in ultrapure water*
8. Pour off water, remove any water traces by rinsing the jar with acetone three times, each time dry the outside of the jar
9. Following the 3^rd^ wash, sonicate the coverslips immersed in acetone for 10 minutes

#### Treat Coverslips with Silane

1. Preheat oven to 110 °C
2. Warm up silane, mPEG-SVA, and biotin-PEG-SVA to room temperature in the vacuum desiccator dome for at least 30 min
3. Mix 125 mL acetone with 2.5 mL of silane in glass beaker, stir using the 5-mL serological pipette
4. Pour off acetone from staining jars, replace with acetone-silane mix, completely submerging coverslips
5. Incubate for 2 min with gentle mixing on a smooth surface (for details, see video in Tanner et al., 2009)
6. Quench reaction over a sink by pouring 1L of ultrapure water into each jar
7. Rinse each jar with ultrapure water 3-4 times
8. Place coverslips on a drying rack with “good side” facing up (the side that faced the solvent in the jar)
9. Bake coverslips at 110 °C for 30 min to cure the silane
10. Place silane bottle in the vacuum dome, flood with argon, store at 4°C in a jar with desiccant for up to one year

#### PEGylate the Coverslips

1. Prepare 0.1 M NaHCO_3_ pH 8.2 by dissolving 430 mg NaHCO_3_ in 50 mL ultrapure water
2. Dissolve 10 mg of biotin-PEG-SVA in 500 μL of 0.1M NaHCO_3_
3. Dissolve 65 mg of mPEG-SVA in the previous solution
4. Place a coverslip with the “good side” facing up on a clean surface, spot 100 μL of PEG solution in the middle, place another coverslip with the “good side” facing down, repeat for a total of 5 such “sandwiches”
5. Place the coverslip sandwiches in a plastic container with a lid, add a wet paper towel in a corner to prevent coverslips from drying, seal the container, incubate at room temperature for 3 hours
6. Place the vials with dry PEG in the vacuum dome, flood with argon, store at −20°C in a jar with desiccant for up to one year
7. Gently peel coverslips apart, thoroughly rinse each one with ultrapure water, keep track of PEGylated side, gently dry with compressed air
8. Place the washed coverslip with the PEG side up in a clean plastic container with a lid, store under vacuum in a desiccation dome for up to 2 months
9. Up to three flow cells can be prepared from each PEGylated coverslip

### Generating Custom Polyclonal Antibodies

To immunodeplete a protein of interest from *Xenopus* egg extract, large amounts of high-quality antibodies are needed (typically 0.2 - 0.4 mg of IgG per KEHRMIT experiment) making the use of commercial antibodies prohibitively expensive. Moreover, immunodepletion-grade antibodies against *Xenopus* replication proteins are usually not available commercially. We routinely generate and validate our own custom rabbit polyclonal antibodies as outlined below (Fig. 10). Animal-free methods such as yeast surface display are an attractive strategy to consider for generating depletion-quality antibodies.

**Figure 10.**
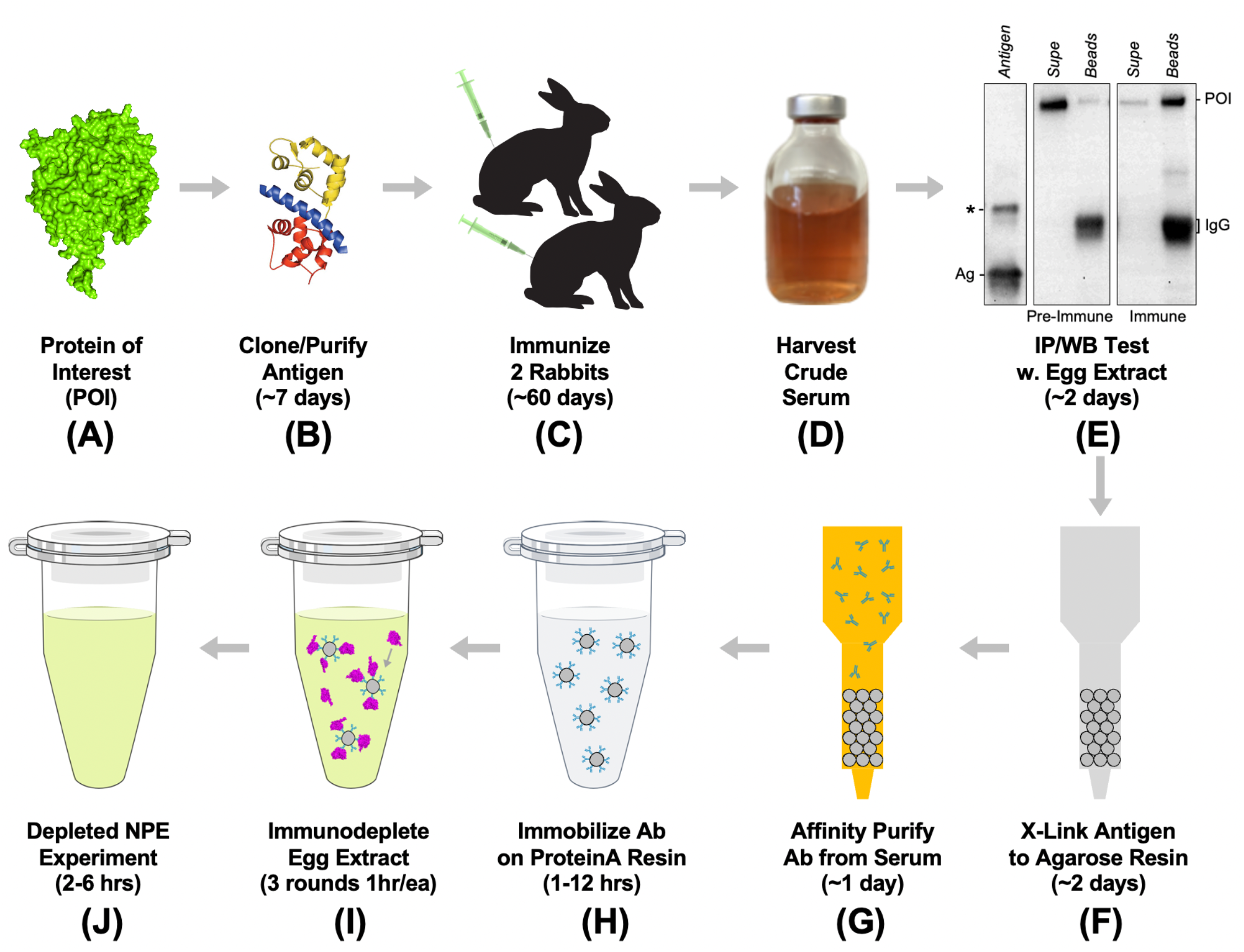
Workflow for Antibody Generation and Validation. **(A-B) The protein of interest is divided into** antigen fragments (each 100-200 amino acids long) that are then expressed in *E. coli* and purified via NiNTA chromatography. **(C)** These antigen fragments are used to immunize rabbits. **(D-E)** Rabbit serum is used for test immunoprecipitations from NPE and HSS. **(F-G)** If the serum contains antibodies that specifically deplete the POI, polyclonal antibodies are affinity purified from the serum. To this end, the antigen fragment is cross-linked to AminoLink resin and incubated with the serum to capture antibodies that bind the antigen. **(H-J)** Affinity-purified antibodies are used to immunodeplete the POI from egg extracts.

#### Selection & Expression of Antigen

1. In an ideal scenario, polyclonal antibodies should be raised against the full-length protein of interest (POI). Most DNA replication and repair proteins are large (>50kDa) and are often part of multi-protein complexes, making the expression and purification of full-length proteins challenging. To simplify the antibody generation pipeline, we select 100-300 amino-acid segments from the POI to be expressed in *E. coli* (Fig. 10A-B). Typically, we clone 3-4 antigen fragments for each POI,which ideally cover the entire polypeptide sequence. To decide where each antigen fragment starts and ends, we use a combination of the following: (a) BepiPred 2.0 – a bioinformatics tool to predict the antigenicity of the polypeptide and identify likely epitopes (Jespersen et al., 2017); (b) any available protein structures or structure prediction tools (like AlphaFold (Jumper et al., 2021) and RoseTTAFold (Baek et al., 2021)) that help identify natural domain boundaries as individual domains are more likely to fold correctly and remain stable when overexpressed; (c) protein disorder prediction tools like IUPred3 and flDPnn to identify likely disordered regions that may be easily accessible to antibodies (Erdos et al., 2021; Hu et al., 2021).
2. Clone the selected 100-300 aa protein fragments from a *Xenopus* cDNA library (prepared from mRNA extracted from oocytes). Alternatively, the DNA sequence could be codon-optimized for expression in bacteria and synthesized commercially as a gene block. We usually clone these constructs into the pET28b(+) vector. Place a 6xHis affinity tag at the beginning or end of the sequence as this tag is compatible with both native and denaturing purification protocols. It is important to leave the natural termini of the protein unmodified as affinity tags can decrease their antigenicity. For example, if the antigen includes amino acids 1-150 from the POI, we would use a C-term 6xHis tag, thus leaving the natural N-terminus intact.
3. Express the antigen in Rosetta DE3 pLysS *E. coli* upon induction with 0.5-1.0 mM IPTG. We usually express antigens for 3 hr at 37 °C or overnight at 16 °C, using 1-2 L of bacterial culture. Bacteria are pelleted, washed with 1x PBS, pelleted again in 50 mL conical tubes, flash-frozen in liquid nitrogen, and stored at −80 °C.

### Antigen Purification

1. On average 50% or more of all antigen fragments that we design express in sufficiently high amounts for antibody generation – i.e. at least 1 milligram of protein from a few liters of bacterial culture. If the antigen fragment is soluble, it can be purified using NiNTA chromatography in native folded conditions using standard protocols (Crowe et al., 1994). Depending on the purity of the NiNTA eluates, additional ion exchange or gel filtration chromatography steps may be needed to eliminate contaminants. However, in many cases the antigen expresses well in bacteria, but is insoluble. In those cases, the antigen is purified under denaturing conditions in the presence of 6 M urea. Since denaturing purification also results in higher protein purity and yield, this has become our preferred method even for soluble or partially soluble proteins. An added benefit of the denaturing purification is that no additional purification steps are needed, and the NiNTA eluate containing 6 M urea and 200-400 mM imidazole can be used directly to immunize rabbits. Finally, in some cases, unfolded antigens may yield better antibodies as all possible epitopes are exposed.
2. Pool elution fractions, concentrate the protein if necessary, prepare aliquots (usually 50-100 μg of antigen each), and ship to the rabbit farm for animal immunizations (our lab uses Cocalico Biologicals). Two male rabbits are injected with the same antigen to compensate for individual animal responses (Fig. 10C). Initial injections (usually 100-200μg of antigen) are administered at day 0 and subsequent boosts (50-100 μg of antigen each) at days 14, 21, 49. Additional boosts (50-100 μg of antigen each) are performed every 28 days after that until sufficient serum has been collected (Fig. 10D). One pre-immune bleed (5-10 mL/animal) is collected on day 0, and two test bleeds (5-10 mL/animal) are collected at days 35 and 56 respectively. Subsequently, production bleeds (15-30 mL/animal) are collected every 14 days.

### Testing Serum

1. We usually test the serum from the second test bleed against the pre-immunization bleed (Fig. 10E). We determine whether (i) the serum can recognize the antigen and the endogenous protein from HSS and NPE on a western blot and (ii) the serum can immunoprecipitate a significant fraction of the POI from HSS or NPE (Fig. 10E).
2. If the serum efficiently immunoprecipitates the POI (preferably without precipitating off-target proteins), we affinity purify antibodies from the serum. To do so 1-5 mg of native or denatured antigen is purified from bacteria using HEPES or phosphate as buffering agents (do not use Tris or other buffers that contain amino groups). The purified protein is dialyzed extensively (3 rounds overnight) to remove any traces of imidazole or other small molecule contaminants. The dialyzed protein is then cross-linked to AminoLink Plus resin (Thermo Fisher #PI20501) following manufacturer recommended protocols (Fig. 10F). If the antigen is insoluble and can only be purified in denaturing conditions, we dialyze the protein extensively into fresh 6M urea at room temperature and perform the crosslinking in that buffer (cross-linking efficiencies range from 50% to 95%). Antibodies that specifically recognize the antigen are then affinity purified from the serum using the standard protocol suggested by the manufacturer (Fig. 10G). Affinity purified antibodies are dialyzed into 1xTris buffered saline (TBS) pH 7.5 (optionally supplemented with 10% sucrose), concentrated to 1.0 mg/mL final concentration, aliquoted, snap-frozen in liquid nitrogen, and stored at −80C for up to several years.

### Purifying Recombinant Proteins

Recombinant proteins for use in KEHRMIT, PhADE, or CIDER can be generated using a variety of recombinant expression systems: *E. coli*, insect cells, mammalian cells, and in vitro transcription-translation (IVTT). We previously expressed recombinant GINS, Cdc45, and CMG in insect cells (see detailed protocols in Low et al., 2020; J. L. Sparks et al., 2019) and recombinant HpaII, Fen1^mKikGR^, and RPA^mKikGR^ in bacteria (see detailed protocols in Loveland et al., 2012; Modesti, 2011; J. L. Sparks et al., 2019). Importantly, the biochemical activity of each recombinant protein should be assessed using in vitro assays. For example, although recombinant GINS can be purified from bacteria, it has a much lower specific biochemical activity that the GINS purified from insect cells (data not shown). Typically we employ a plasmid replication assay where the protein of interest is immunodepleted from extract and the depleted extract is supplemented with the recombinant protein at a physiological concentration (Lebofsky et al., 2009; Sparks and Chistol et al., 2019).

### Fluorescent Labeling of Recombinant Proteins

To visualize a recombinant protein, the protein must first be labeled with a fluorophore. To correlate the fluorescent intensity of a diffraction-limited spot on the detector to the number of proteins within that area, the protein of interest must be labeled with a single dye. Therefore, non-specific labeling methods such as the NHS-ester for attachment to primary amines are not recommended. Although protein fusions to fluorescent proteins achieve this stoichiometric labeling, labeling with organic dyes remains preferable for two reasons. First, organic fluorophores have superior photophysical characteristics (extinction coefficient, quantum yield, and photostability). Second, organic fluorophores are much smaller (~1 kDa) than fluorescent proteins like GFP (~26 kDa) and therefore less likely to interfere with the biological function of the POI. For example, while Cdc45^AF647^ (labeled via the Sortase approach, see below) efficiently rescued replication in Cdc45-depleted extract, Cdc45 tagged with SNAP at either terminus did not, suggesting that the ~20kDa SNAP tag impairs the function of Cdc45. Of course, some proteins may tolerate bulky fusion proteins, but this must be tested experimentally.

### Enzymatic Labeling of Peptide Tags

To date, several enzymatic modifications to sequence-specific protein tags have been established as robust tools for labeling proteins. Although many different tools are commonly used (BirA, LplA, TTL, Sfp/AcpS, TGase, AnkX/Lem3, SrtA, FGE), they all follow the same general workflow (Lotze et al., 2016). Recombinant protein is prepared with a genetically encoded short peptide tag (typically 5-20 aa long) and subsequently incubated with both the labeling enzyme and the fluorescent substrate. The substrate is usually a small molecule or a peptide conjugated to an organic dye like Cy3/5 or AlexaFluor488/546/647. Fig. 11A presents an an overview of the method we commonly use – SortaseA mediated peptide conjugation (Antos et al., 2017; Guimaraes et al., 2013; Popp et al., 2007).

**Figure 11.**
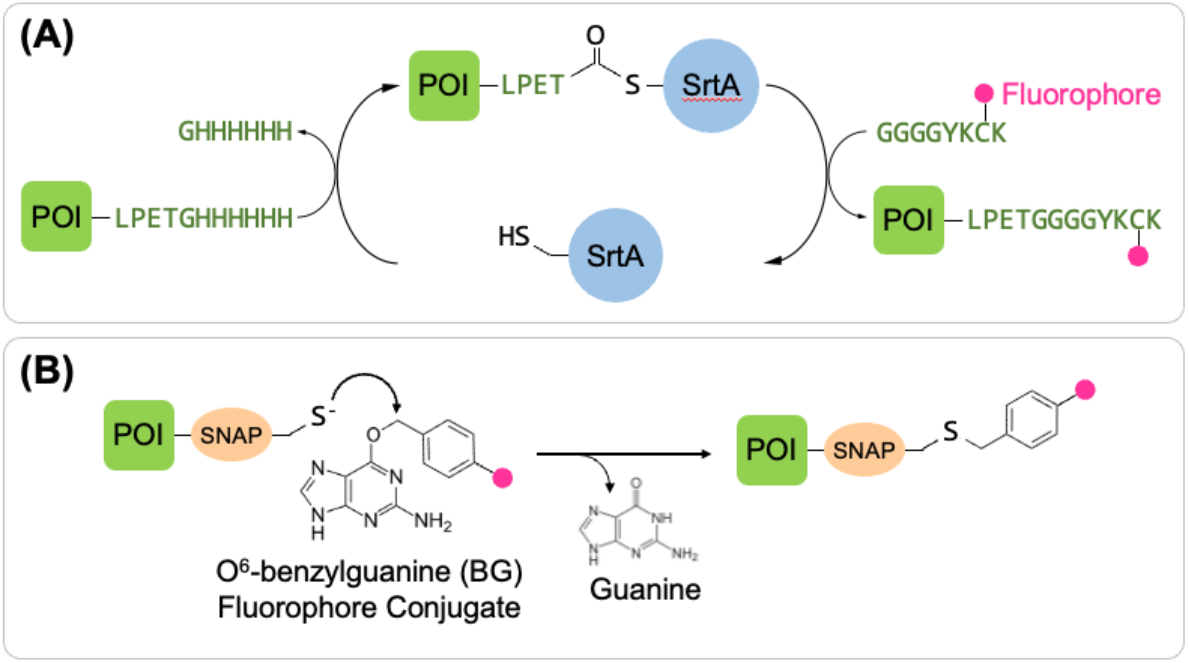
Overview of SortaseA-Mediated and SNAP-Mediated Labeling of Recombinant Proteins.

### Self-Modifying Enzymes

Another attractive method for site-specific stoichiometric protein labeling is the use self-modifying enzymes. Three are commercially available orthogonal self-labeling approaches: SNAP-tag (NEB), CLIP-tag (NEB), and HaloTag (Promega) (Wilhelm et al., 2021). The SNAP-tag is derived from the 20 kDa DNA repair protein O^6^-alkylguanine-DNA alkyltransferase and has been engineered to instead react specifically with benzylguanine (BG) derivatives (Cole, 2013). The CLIP-tag is a modified SNAP-tag such that its substrate specificity is different, reacting instead with the substrate O2-benzylcytosine. In practice, both the SNAP-tag and CLIP-tag can be used in the same reaction as their substrate specificities are very high (Hoehnel & Lutolf, 2015). BG substrates conjugated to AlexaFluor dyes ranging from 488 to 647 nm are commercially available from NEB. The 33 kDa HaloTag® was derived from the enzyme haloalkane dehalogenase and reacts with chloroalkanes (Los et al., 2008). HaloTag substrates conjugated to Janelia Fluor® dyes ranging from 525 to 650 nm are available from Promega. Fig. 11B summarized the SNAP-tag labeling workflow – another method we commonly use.

### Validating Recombinant Proteins and Affinity Purified Antibodies via Immunodepletion-Rescue Assay

Before antibodies and fluorescently labeled recombinant proteins can be used in single molecule experiments, they must be first validated via a robust and reproducible biochemical assay (see Table 2). Specifically, we verify that the POI is efficiently immunodepleted from extract via western blot, and that the depletion of this protein impairs or completely abolishes DNA replication. If the addition of recombinant protein rescues this defect, we are confident that the immunodepletion was specific and no other essential factors were depleted non-specifically. If the fluorescently labeled recombinant protein also robustly rescues the immunodepletion, it indicates that the fluorescent tag does not interfere with protein function. Below we present a detailed protocol for the depletion-rescue biochemical assay for GINS – the protein used to visualize the CMG helicase in the original version of the KEHRMIT assay (Sparks and Chistol et al., 2019). The pipeline for purifying recombinant proteins and validating custom antibodies is summarized in Fig. 12. For additional pointers related to the biochemical DNA replication assay, pls refer to the detailed protocols published in Lebofsky et al., 2009.

**Table 2:**
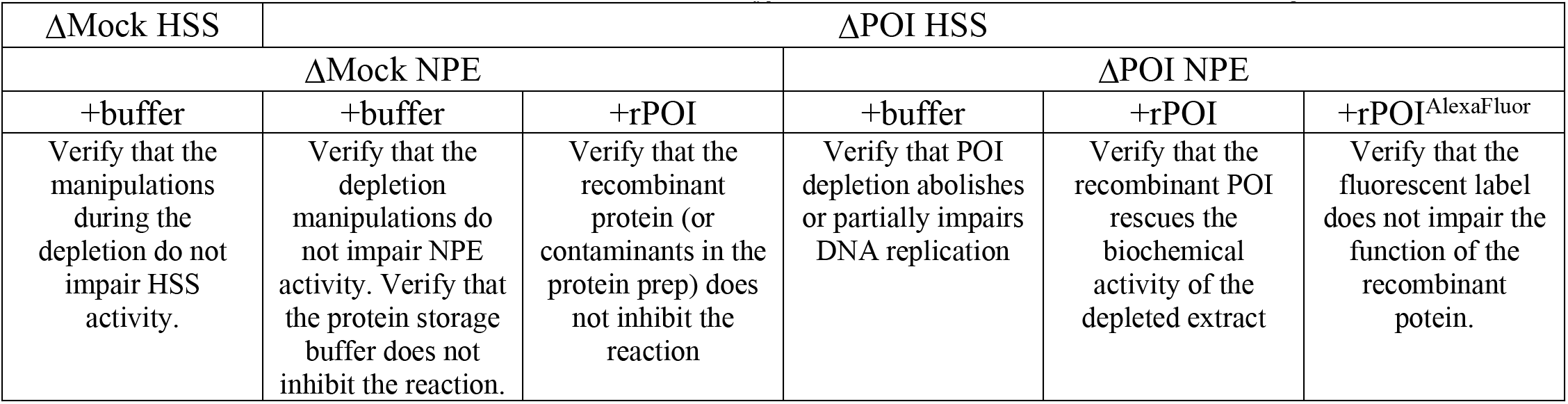
Scheme for Testing the Function of Recombinant Proteins via Depletion-Addback Experiments. To rule out non-specific depletion effects, the controls listed here must be performed in a biochemical DNA replication assay. Briefly, the DNA is licensed in either mock-depleted or POI-depleted HSS, then the replication reaction is initiated by mixing licensed DNA with NPE. The NPE (mock-depleted or POI-depleted) is supplemented with POI storage buffer, recombinant POI, or recombinant POI that has been fluorescently labeled.. The notes below each reaction condition clarify what each individual control accomplishes.

**Figure 12.**
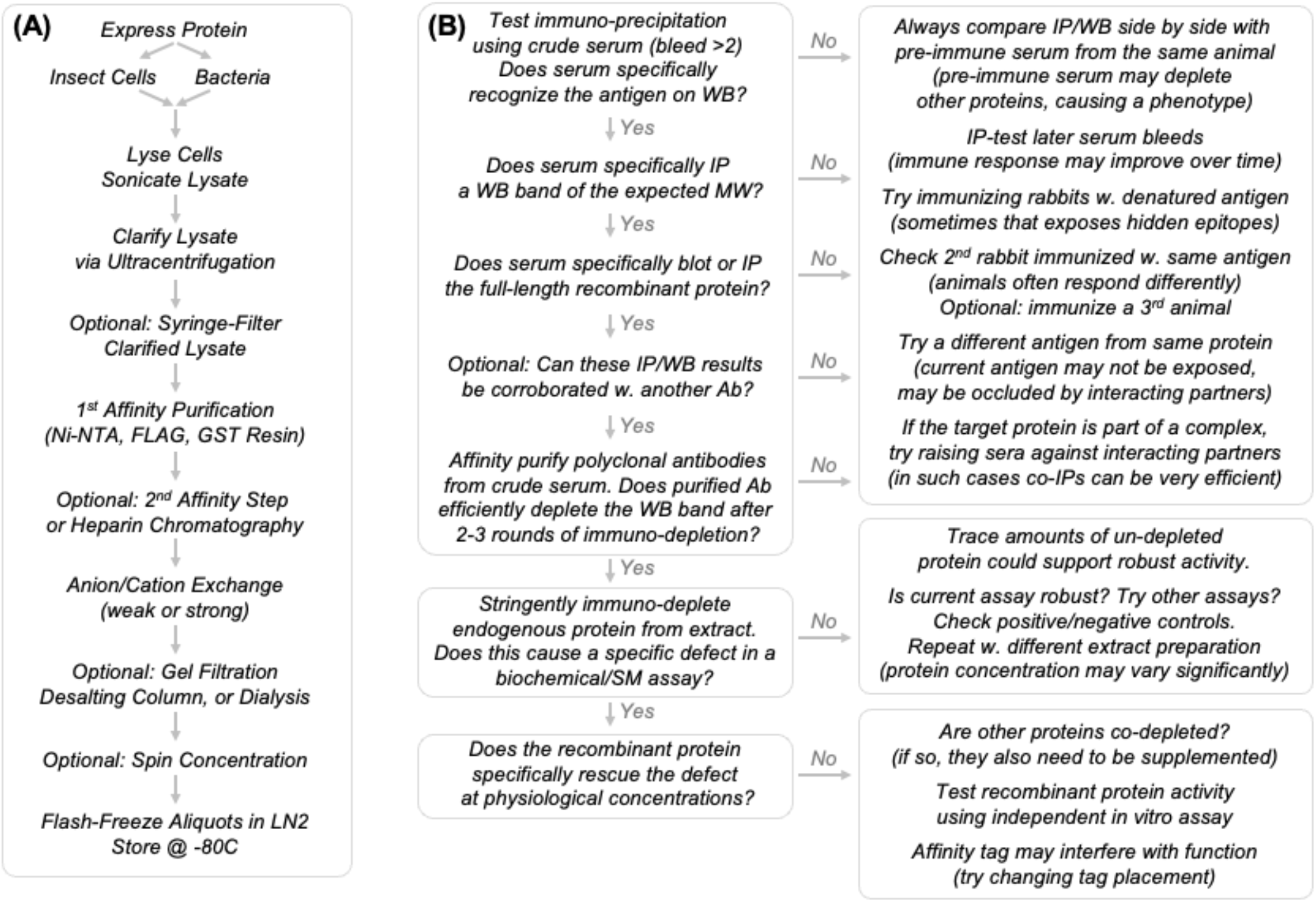
Workflow for Purifying Recombinant Proteins and Validating Custom Antibodies. **(A)** Summary of our strategy for purifying any recombinant protein for use in KEHRMIT experiments. **(B)** Summary of our strategy for testing custom antibodies raised against the protein of interest. We list several troubleshooting steps in this process.

**Figure 13.**
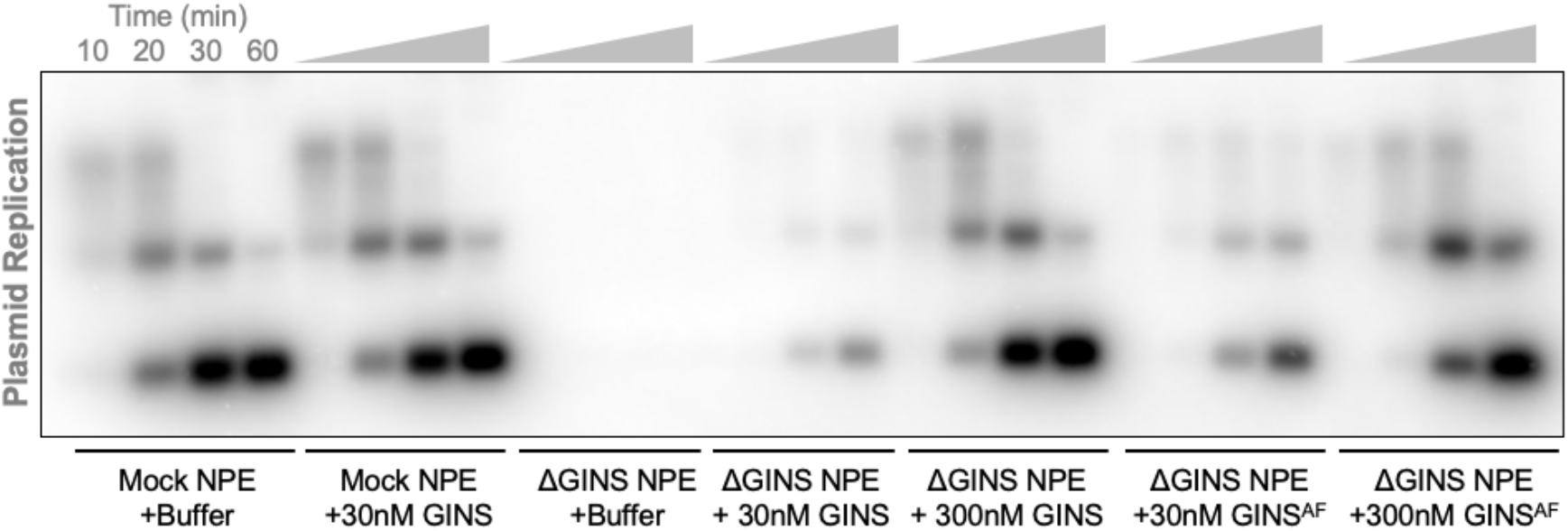
Example of a Biochemical Assay Designed to Test the Activity of Recombinant GINS. All conditions used GINS-depleted HSS that we previously showed to retain robust licensing activity (GINS is not involved in DNA licensing). In all cases 99% or more of the endogenous GINS was immunodepleted (assessed via Western Blotting). This experiment reveals the following (from left to right): (1) the protein storage buffer does not impair replication; (2) the recombinant GINS itself does not impair replication; (3) depleting GINS abolishes DNA replication; (4) small amounts of recombinant GINS partially rescue replication in depleted extract; (5) physiological concentrations of recombinant GINS (~300 nM) fully rescue replication; and (6) the fluorescent labeling of recombinant GINS with Alexa Fluor dyes does not interfere with the biochemical activity of this protein.

#### Extract Preparation

The protocol below is meant to illustrate the antibody and protein validation workflow. For technical details on performing depletions see the detailed KEHRMIT protocol.

1. Prepare two samples of HSS (cytoplasmic extract that recapitulates G1-phase)
  a. Optional: HSS mock-depleted with non-specific IgGs (purified from pre-immunization serum via ProteinA-S epharose chromatography).
  b. HSS depleted with anti-GINS antibodies affinity purified from post-immunization serum.
2. Prepare two samples of NPE (nuclear extract that recapitulates S-phase):
  a. NPE mock-depleted with non-specific IgGs.
  b. NPE depleted anti-GINS antibodies.
3. License a 3-5 kbp plasmid (we usually use pBlueScript) in HSS supplemented with ATP regeneration system (ARS) and radioactively labeled dNTPs for monitoring DNA synthesis. Each licensing reaction contains the following (the volume may be scaled up or down depending on the specific experiment):
  ○ 20 μL GINS-depleted or mock-depleted HSS (for depletion details see KEHRMIT protocol below)
  ○ 1μL pBlueScript plasmid at 180 ng/uL (7.5ng/μL f.c.)
  ○ 1 μL ARS (for details see the KEHRMIT protocol below)
  ○ 2 μL dCTP [a-^32^P] (PerkinElmer #BLU513A250UC) *Mix well by gently pipetting up-down 20x* *Incubate for 30 min at room temperature*
4. Prepare 50% NPE dilutions (it was previously empirically determined that diluting NPE to 50% maximizes its DNA replication activity, but this must be tested for each extract preparation)
5. Separate 50% NPE mixes must be prepared for each condition. These are used to validate various aspects of antibody and recombinant protein preparations, as summarized in Table 2.

a. mock-depleted NPE + buffer
b. mock-depleted NPE + rGINS
c. mock-depleted NPE + rGINS^AF647^
d. GINS-depleted NPE + buffer
e. GINS-depleted NPE + rGINS
f. GINS-depleted NPE + rGINS^AF647^
6. Each individual reaction contains the following (can be scaled up or down to minimize extract waste):

○ 5 μL mock-depleted or GINS-depleted NPE
○ 0.5-1.0 μL protein storage buffer or recombinant protein (no more than 10% of the reaction volume)
○ 0.5 μL ARS
○ 3.5-4.0 μL ELB-Sucrose (to make up a final volume of 10 μL) *Mix well by gently pipetting up-down 20x* *Incubate for 10 min at room temperature*
7. Set up individual replication reactions by mixing DNA licensed in HSS with 50% NPE:

a. ○ 3 μL of licensing reaction
b. ○ 6 μL of 50% NPE mix *Mix well by gently pipetting up-down 20x*
8. At specific time points (typically 10, 20, 30, 60 min) remove 2.0 μL of the reaction and mix with 15 μL of Replication Stop buffer (8 mM EGTA, 0.13% phosphoric acid, 10% Ficoll, 5% SDS, 0.2% Bromophenol blue, 80 mM Tris at pH 8.0).
9. To each time point sample add 0.5 μL of Proteinase K (NEB #P8107S) and incubate for 30 min @ 37 °C to digest all the proteins.
10. Run the reaction on a 0.8% agarose gel in 1xTAE or 1xTBE, cut the gel along the bromophenol blue migration front, discard the bottom portion of the gel containing excess radioactive dNTPs, keep the top portion of the gel that contains the plasmid replication products and intermediates.
11. Sandwich the gel between two sheets of DNA/RNA transfer membrane (Pall BiodyneB 0.45μm #60207). Sandwich the membrane-gel-membrane assembly between two sheets of Whatman paper (GE Healthcare #3030-917). Dry this assembly between paper towels for 1hr, then on a gel drier for 1-2 hrs.
12. Wrap the dried gel in cling plastic (grocery store grade) and expose on a phosphorscreen for ~2hrs. Image the phosphorscreen on a Typhoon imager (GE Healthcare).

### Microscope Configuration and Imaging Settings

Below we describe the TIRF microscope configuration (Fig. 14) needed to conduct KEHRMIT, PhADE, and CIDER experiments. The list of equipment should include the following:

○ Microscope body and motorized XY stage, preferably with encoders for micron precision
○ 100X NA1.49 apochromat infinity-corrected oil immersion objective, preferably optimized for TIRF
○ Laser launch with 405nm, 488nm, 532nm or 561nm, 640nm lasers, (>1mW/ea out of the objective)
○ TIRF illuminator, preferably with motorized control of the TIRF angle
○ Temperature-controlled microscope enclosure to minimize drift
○ Blackout panels or curtains to minimize external light pollution
○ Syringe pump for precise control of buffer and extract flow rates and volumes
○ Laser line rejection dichroic and assorted emission filters
○ Automated focus lock system to compensate focus drift over the course of 10-100min experiments
○ Camera(s) suitable for single fluorophore detection (as discussed below)

**Figure 14.**
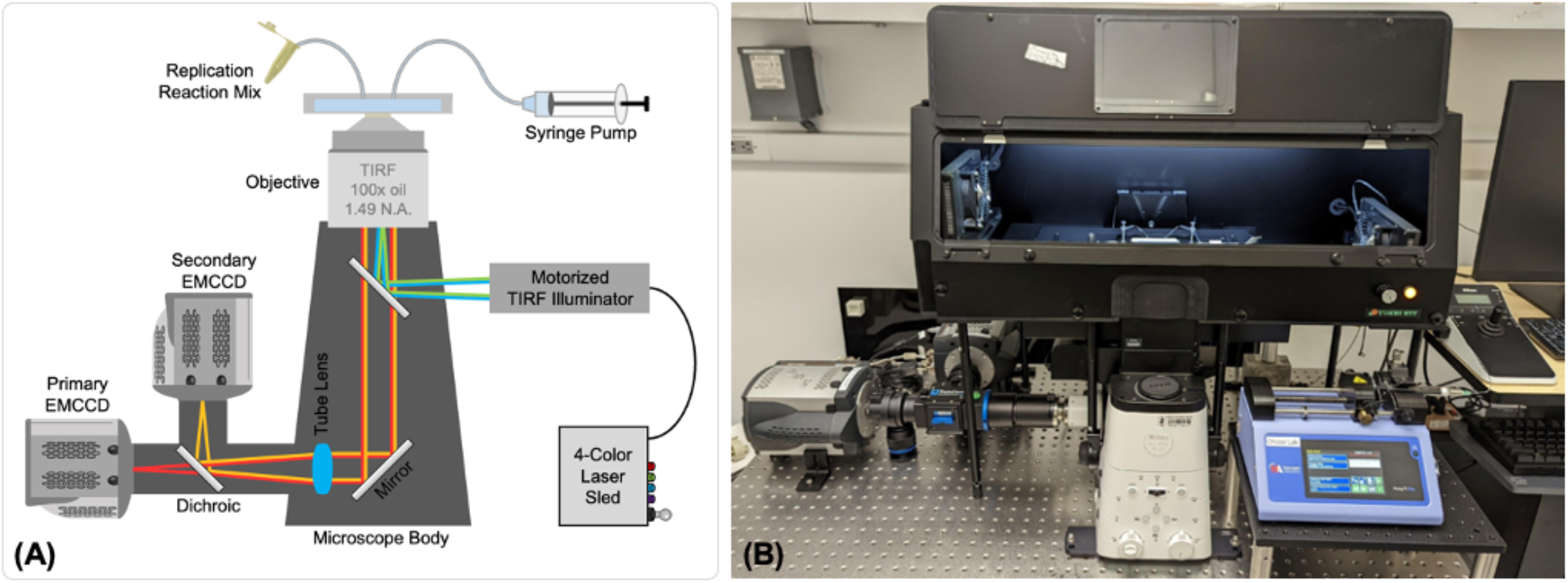
Overview of our Microscope Used for KEHRMIT Experiments. **(A)** Schematic diagram highlighting the key components (not to scale). **(B)** Photograph of our instrument provided for clarity. Note the temperature control enclosure that doubles as a dark box.

#### Microscope

We use a state-of-the-art commercial Nikon TIRF microscope built around the Ti2 body with a motorized XYZ stage and Perfect Focus system (Fig. 14). However, KEHRMIT experiments can be performed on any commercial or home-built microscope with a motorized stage and a reliable focus lock system. The latter is critical for high-throughput imaging where several fields of view are periodically imaged over the course of 10-100 min. Without an autofocus system, it should be possible to image a single field of view, but the focal plane may drift significantly over the course of several minutes. Another important microscope accessory is the temperature control enclosure (Tokai Thermobox) that plays three key roles: (i) provides temperature control and day-to-day consistency for the biochemical reaction of DNA replication (24 °C results in the fastest replication forks); (ii) provides temperature control for the instrument, minimizing objective thermal expansion and focus drift; (iii) our enclosure is made of black plastic and acts as a blackout box, enabling us to carry out single-molecule experiments even when the lights are on in the room.

#### Focusing

To maximize the signal-to-noise ratio it is critical to focus on the double-tethered DNA molecules, which are above the coverslip surface (Fig. 2A). The simplest method is to lock the autofocus system on the SYTOX-stained tethered DNA before any extract is introduced into the flow cell. However, there is a chance that the focal plane may drift when flowing buffers or extract into the flow cell, perhaps due to accidentally touching the motorized stage. The second method is to mix trace amounts of streptavidin^AlexaFluor647^ with unlabeled streptavidin during surface functionalization at the start of the experiment. This adds fiducial markers (preferably 10-50 spots per field of view) to the surface of the coverslip, and since they are fixed, they will not be mistaken for replisomes. Separately, we measure the Z-offset between the surface-bound fluorescent streptavidin markers and the plane in which tethered DNA is in focus – in our experiments this offset is 0.3-0.4 microns of Z-axis piezo movement (this offset does not change between experiments). Immediately before beginning real-time KEHRMIT imaging, we focus on the surface fiducial markers, enter the known Z-offset, and set the focus lock system to maintain that focal plane. Although more convoluted than the first method, this strategy provides more consistent results.

#### Camera(s)

Our microscope is configured with two Andor iXon 897 Ultra EMCCDs (Oxford Instruments) for simultaneous full-chip two-color imaging via a TwinCam image splitter (Cairn Research). This enables high temporal resolution imaging and improved mechanical stability (as there is no need to switch filters with a motorized wheel to achieve two-color imaging) but comes at increased cost and complexity. Importantly, for each experiment the two-color channels must be aligned using a slide with multi-spectral fluorescent beads (0.1 micron TetraSpeck beads, Thermo Fisher #T7279). Precise alignment is then performed computationally by using reference images of the bead slide in the color channels. To enable 3-color and 4-color imaging, we use a single camera and the motorized filter wheel built into the microscope, but this comes at the cost of reduced throughput (fewer fields of view per experiment) or lower time resolution (fewer frames per minute), as individual color images must be acquired sequentially. Finally, although the performance of CMOS cameras has improved dramatically over the past decade, EMCCDs still outperform CMOS cameras for ultra-low light imaging (which we employ to minimize photobleaching).

#### Microfluidics

A microfluidic system is needed to deliver buffers and extract into the flow cell where the biochemical reaction takes place in a KEHRMIT experiment. This system must be able to precisely pump volumes as little as 10 μL at rates as low as 10 μL/min. To this end, we use the Programmable Syringe Pump 11 Elite (Harvard Apparatus #70-4504) in “withdraw mode” coupled with a 5 mL gastight glass syringe (Hamilton #1005). This pump-syringe combination has an error of ~1 μL when withdrawing volumes of 10-500 μL.

It is critical that tiny air bubbles are prevented from forming inside the microfluidic flow cell, as these bubbles break double-tethered DNA molecules, interfere with buffer-extract exchange, and completely disrupt focus locking. To minimize the chances of air bubbles forming and remaining trapped in the flow path, all buffers are thoroughly degassed for ~30min before the experiment. During the initial phase of setting up the single-molecule experiment, a large volume of buffer is flown rapidly through the flow cell while the outlet tubing is flicked vigorously to generate hydrostatic shock and dislocate any microscopic bubbles from the flow cell and tubing (see details in the DNA tethering protocol below). Importantly, the buffer is supplemented with detergent to lower the surface tension of water and disrupt bubbles.

#### Optimizing Imaging Parameters

When performing single-molecule imaging experiments at relatively high concentrations of fluorescent proteins in solution (>30nM), where it is difficult to obtain a high SNR (>2), a few factors should be considered to improve data quality.

a. The TIRF angle influences both the depth of the evanescent illumination field and the intensity of the field at all depths. Higher TIRF angles result in a shallower evanescent field and lower light intensities. Since the fluorescent proteins are not attached directly to the coverslip, but are bound to double-tethered DNAs that fluctuate above the glass coverslip, an intermediate TIRF angle should be selected. Our microscope is equipped with a motorized TIRF illuminator that provides precise control of the illumination angle, and reproducibility for experiments over several months.
b. The laser power directly affects the number of photons collected from each fluorophore during each exposure. When the number of photons collected per pixel is above 100, the EMCCD camera is no longer the limiting factor for SNR (consult the Andor iXon 897 manual for details on how to measure the number of photons collected). For experiments with high concentrations of fluorescent protein, the best SNR is achieved by increasing the exposure duration and decreasing the laser power (which also minimizes photobleaching), and keeping the number of photons per pixel below ~500. Longer exposures (500-1000 ms) will average the background signal due to freely diffusing fluorophores, helping to improve SNR.

### Detailed Protocol for a KEHRMIT Experiment

#### Prepare Reaction Buffers

10X ELB Salts:

○ 100 mM HEPES
○ 500 mM KCl
○ 25 mM MgCl_2_ *pH to 7.7 with KOH sterifilter, wrap in foil store at 4 °C for up to 1 yr*

1xELB-Sucrose:

○ 1X ELB Salts
○ 0.25 M Sucrose *sterifilter, aliquot store at 4 °C for up to 1 month degas for 30 min prior to using*

DNA Buffer:

○ 20 mM Tris-HCl pH 8.0
○ 50 mM NaCl
○ 0.5 mM EDTA *sterifilter, aliquot store at 4 °C for up to 1 month degas for 30 min prior to using*

#### Prepare ATP Regeneration System Components

Adenosine Triphosphate (ATP): 0.2 M ATP (Sigma #A7699) in ultrapure water *pH to 7.0 with NaOH sterifilter, make 20 °L aliquots store at −20°C for up to 1 yr*
Phosphocreatine (PC): 1.0 M PC (Sigma #P6502) in 20 mM KH_2_PO_4_ *pH 7.0 sterifilter prepare 40 μL aliquots store at −20°C for up to 1 yr*
Creatine Phosphokinase (CPK): 5 mg/mL CPK (Sigma #C3755) in 20 mM HEPES pH 7.5, 50% glycerol, 50 mM NaCl *prepare 50 μL aliquots store at −20°C for up to 1 yr*

#### Binding Antibodies to ProteinA Sepharose Beads

For a standard KEHRMIT experiment 50 μL of NPE and 70 μL of HSS must be depleted of GINS (or other POI). To this end, anti-GINS antibodies and mock IgGs must first be bound to ProteinA Sepharose (rProtein A Sepharose Fast Flow #GE17-1279-03). Typically, three rounds of immunodepletion are performed for each extract, but depending on the quality of the antibodies and the concentration of the POI, two rounds of immunodepletion may be sufficient (or in the worst-case scenario – 4 rounds may be necessary). This is determined by extensive testing and validation of the antibody. Finally, for each 1 volume of extract, 1/5 volume of PAS beads (“dry” volume) must be used to ensure efficient mixing during the depletion. The amount of antibody necessary to efficiently deplete the POI varies, but a good starting point is to use 1 volume of anti-POI antibody at 1mg/mL for every 1 volume of extract. This ratio is empirically tested, and the amount of antibody may be increased or decreased by up to 2x from that initial starting condition. Note that the PAS beads are stored at 4°C as a 33% slurry in 1xPBS supplemented with sodium azide, so every 1 μL of PAS beads (dry volume) is equivalent to 3 μL of 33% slurry. Since PAS beads settle quickly the PAS slurry must be thoroughly resuspended by inverting the tube and pipette mixing immediately before pipetting.

1. <u>Calculate the amounts of PAS and antibody needed for the immunodepletion</u> To deplete 50 μL of NPE: (10 μL of PAS beads + 50 μL of anti-GINS antibody at 1mg/mL) * 3 rounds To deplete 70 μL of HSS: (14 μL of PAS beads + 70 μL of anti-GINS antibody at 1mg/mL) * 3 rounds Total amounts: 72 μL of PAS beads (216 μL of 33% PAS slurry) + 360 μL of anti-GINS antibody at 1 mg/mL
2. <u>Wash PAS Beads Twice with 1x ELB Sucrose Buffer</u>
  ○ Pipet 216 μL of 33% PAS slurry into a siliconized 0.65mL Eppendorf tube (Corning #3206)
  ○ Pellet beads via centrifugation for 30 sec at 2000 g (preferably in a swing-bucket rotor)
  ○ Aspirate the supernatant with an ultra-fine gel loading tip (Fisherbrand #02-707-88) *(the aspiration tip is connected via tygon tubing to house vacuum via a vacuum trap)*
  ○ Add 400 μL of 1xELB sucrose, resuspend beads
  ○ Pellet beads as above, aspirate supernatant (ultrafine tip can be reused several times)
  ○ Repeat the wash with 400 μL of 1xELB sucrose,
  ○ Pellet beads, aspirate supernatant (beads should not be kept “dry” for longer than ~30sec)
3. <u>Incubate Washed PAS Beads with Anti-GINS Antibody</u>
  ○ To the “dry” beads add 360 μL of anti-GINS affinity purified antibody at 1 mg/mL
  ○ Resuspend beads by gentle pipette mixing
  ○ Incubate on a rotator (Southwest Science #STR200-V) overnight @ 4C (or at RT for at least 1 hour)
4. <u>Wash the PAS Beads to Remove Unbound Antibody</u>
  ○ Pellet beads via centrifugation for 30 sec at 2000g
  ○ Optional: verify concentration of unbound antibody in the supernatant via NanoDrop (should be negligible)
  ○ Aspirate supernatant as indicated above
  ○ Wash the beads twice with 400 μL of 1x ELB sucrose (as indicated above)
  ○ Wash the beads twice with 400 μL of 1xELB sucrose + 500mM NaCl *(stringent high-salt wash to remove non-specifically bound proteins and contaminants)*
  ○ Wash the beads twice with 400 μL of 1x ELB sucrose (in preparation for use with extracts)
  ○ Pellet beads, aspirate supernatant
  ○ Add 300 μL of 1x ELB sucrose and gently pipette mix to prepare a ~20% PAS slurry, aliquot as follows
  ○ Prepare 3 aliquots of 70 μL of 20% PAS in 0.65mL siliconized tubes for HSS depletions (label accordingly)
  ○ Prepare 3 aliquots of 50 μL of 20% PAS in 0.65mL siliconized tubes for NPE depletions (label accordingly)
  ○ There will be a small amount (<10 μL) of PAS slurry left over – discard
  ○ Keep the PAS slurry aliquots on ice

### Immunodepleting GINS from Egg Extracts

Immediately before starting the immunodepletion procedure, extracts must be thawed, and any insoluble material should be pelleted via centrifugation. Only HSS is supplemented with nocodazole to disrupt microtubule filaments (NPE already contains nocodazole). Pipet gently to avoid introducing small air bubbles in the extract, avoid “bubble foam” at all costs. Extracts should be kept cold for the entire duration of the immunodepletion. Each round of immunodepletion requires a 45 min incubation of antibody-coated PAS beads with extract. Additional handling between depletion rounds may take 5-10 min. Incubations are performed at 4°C in a cold room or a deli fridge with a clear glass door. A single “mixing bubble” (~1 mm in diameter) is intentionally added to the top of the extract-bead slurry before beginning the 45-min incubation – this helps mix the viscous extract-bead slurry during end-over-end rotation-incubation. Note that good mixing of the slurry during this step is critical for efficient immunodepletions.

1. <u>Clarify HSS Prior to Immunodepletion</u>
  ○ Thaw 80 μL of HSS (2 aliquots of 40 μL/ea)
  ○ Combine extract into a single 0.2 mL PCR tube
  ○ Add 1 μL Nocodazole (0.5 mg/mL in DMSO)
  ○ Pipette mix gently 20x
  ○ Pellet insoluble material via centrifugation for 5min at 15000g at 4°C
  ○ Use only the top 70 μL of extract for depletions, avoiding the insoluble material at the bottom
2. <u>Clarify NPE Prior to Immunodepletion</u>
  ○ Thaw 55 μL of NPE (5 aliquots of 11 μL/ea)
  ○ Combine extract into a single 0.2 mL PCR tube
  ○ Pipette mix gently 20x
  ○ Pellet insoluble material via centrifugation for 5min at 15000g at 4°C (spin at the same time as HSS)
  ○ Use only the top 50 μL of extract for depletions, avoiding the insoluble material at the bottom
3. <u>Set Up the 1^st^ Depletion Round for HSS</u>
  ○ Take a tube with beads reserved for HSS depletions (70 μL of 20% PAS slurry = 14 μL of dry beads)
  ○ Pellet beads via centrifugation for 30 sec at 2000g
  ○ Use an ultrafine tip to aspirate all 1xELB-sucrose buffer until the beads appear dry
  ○ Add 70 μL of clarified HSS onto the beads, resuspend PAS by gently pipette-mixing 10x
  ○ Add a small mixing bubble to the top, incubate on rotator at 4°C for 45 min
4. <u>Set Up the 1^st^ Depletion Round for NPE (same as for HSS, except the following)</u>
  ○ Use tube with beads reserved for NPE depletions (50 μL of 20% PAS slurry = 10 μL of dry beads)
  ○ Use 50 μL of clarified NPE
5. <u>Set Up the 2^nd^ Depletion Round</u>
  ○ Pellet slurry of antibody-coated PAS beads via centrifugation for 30 sec at 2000g at 4°C
  ○ Aspirate all the 1xELB-sucrose buffer leaving beads dry (do not keep beads dry for more than ~30 sec)
  ○ Centrifuge the tube with bead-extract mixture from the 1^st^ depletion round (30 sec at 2000g at 4°C)
  ○ Use a P20 pipette to collect as much clear extract without picking up beads
  ○ Transfer this clear extract to the dry beads reserved for the 2^nd^ depletion round
  ○ Harvest the last few μL of extract using a P10 pipette, insert the tip to the bottom of the bead+extract tube forming a seal between the tube and the tip then slowly tilt the tip to break the seal and gently withdraw the extract (this will minimize bead carryover)
  ○ Gently pipette-mix 10x the beads+extract for the 2^nd^ depletion round
  ○ Add a small mixing bubble to the top, incubate on rotator at 4°C for 45 min
6. <u>Set Up the 3^rd^ Depletion Round</u>
  ○ Same as the 2^nd^ round of depletion
7. <u>Harvest Depleted Extract</u>
  ○ Harvest the extract as described above into a clean 0.65mL tube (do this separately for HSS and NPE)
  ○ Pellet any leftover beads via centrifugation for 30 sec at 2000g at 4°C, keep on ice until needed
  ○ Carefully harvest extract off the top to avoid bead carryover into the single-molecule reaction

### Preparing Licensing, Replication Initiation, and Replication Elongation Mixes

Once the DNA templates are tethered in the microfluidic flow cell (which should already be mounted onto the microscope stage), prepare the three reaction mixes necessary for a KEHRMIT experiment. First, DNA is licensed by flowing the licensing mix into the flow cell and incubating for a few minutes. Second, replication is initiated for a few minutes by incubating the flow cell with the initiation mix containing fluorescently labeled GINS. Finally, free GINS^AF647^ is washed off by flowing replication elongation mix. This prevents any further origins from firing, but the origins that already fired using fluorescent GINS are allowed to replicate DNA for 30-120 minutes (depending on the exact design of the experiment).

Single-molecule replication reactions require the use of carrier DNA for efficient licensing and replication as discussed in detail in previous studies (Lebofsky et al., 2011; Loveland et al., 2012). Importantly, we use a short 30-bp double-stranded DNA fragment as carrier DNA during the licensing reaction as this DNA is too short to stably load Mcm2-7 double hexamers, and therefore this DNA does not get replicated later. For the replication initiation and replication elongation reactions we use the pBlueScript plasmid (~3kb) as carrier DNA. Since this plasmid is incubated in a mixture of HSS and NPE, it does not get licensed, and therefore does not get replicated. Finally, it is critical that all three reaction mixes be allowed to equilibrate to room temperature (RT) in a water bath or a heat block prior to beginning the single-molecule experiment.

1. <u>Prepare ATP Regeneration System (ARS) Mix</u>
  ○ 5 μL ATP (0.2 M)
  ○ 0.5 μL CPK (5mg/mL)
  ○ 10 μL PC (1 M)
  ○ Gently pipette-mix 10x, store at room temperature for up to 30 min
2. <u>Prepare HSS+NPE Mix for Replication Initiation/Elongation Mixes</u> NPE contains a high concentration of geminin that binds to Cdt1 in HSS, rendering this extract mixture non-competent for DNA licensing which requires Cdt1 (Pozo & Cook, 2016). This is done to prevent licensing of carrier plasmid DNA in the replication initiation and elongation mixes.

○ Take 31 μL of GINS-depleted HSS, avoid carrying over trace amounts of PAS beads
○ Take 31 μL of GINS-depleted NPE, avoid carrying over trace amounts of PAS beads
○ Gently pipette-mix 20x, incubate at room temperature for 10 min
3. <u>Prepare DNA Licensing Mix (22 μL total)</u>
  ○ 2.0 μL 30-bp carrier DNA oligo (300 ng/μL)
  ○ 0.5 μL ATP Regeneration Mix
  ○ 20.0 μL GINS-depleted HSS
  ○ Gently pipette-mix 20x, incubate at room temperature for 10 min
4. <u>Prepare Replication Initiation Mix (30 μL total)</u>
  ○ 1.0 μL pBlueScript carrier plasmid DNA (600 ng/μL)
  ○ 1.0 μL ATP Regeneration Mix
  ○ 7.0 μL 1xELB-Sucrose
  ○ 20.0 μL GINS-depleted HSS+NPE mix
  ○ 1.0 μL GINS^AF647^ (3 - 5 μM stock)
  ○ Gently pipette-mix 20x, incubate at room temperature for 10 min
5. <u>Prepare Replication Elongation Mix (60 μL total)</u>
  ○ 2.0 μL pBlueScript carrier plasmid DNA (600 ng/μL)
  ○ 2.0 μL ATP Regeneration Mix
  ○ 15.0 μL 1xELB-Sucrose
  ○ 40.0 μL GINS-depleted HSS+NPE mix
  ○ 1.0 μL Fen1^mKikGR^ (60 μM stock)
  ○ Gently pipette-mix 20x, incubate at room temperature for 10 min

### Constructing the Microfluidic Reaction Chamber

We usually assemble the flow cell during the 1^st^ depletion round, but flow cells may also be prepared several hours in advance and stored under vacuum. The glass slide with pre-drilled holes can be reused several times by incubating old flow cells in acetone overnight, then cleaning epoxy residue with a razor and paper towels dipped in acetone. Flow cell assembly steps are illustrated in Fig. 15.

1. Drill two holes 8 mm apart in the center of a 75 mm x 25 mm glass slide (VWR #16004-422) using a Dremel 4000 rotary tool mounted in a drill press (Fig. 15A). We use a 1-mm diamond dental bur (A&M Instruments #HP863-10-Coarse) to drill the glass. The slide is held in a shallow water bath to capture glass chips and dissipate heat. One hole should snugly fit PE20 tubing (BD Intramedic #427406), and the second hold should fit PE60 tubing (BD Intramedic #427416) (Fig. 15B).
2. Using a sharp scalpel, cut a 4.0 cm segment of PE60 tubing (outlet) and a 5.5 cm segment of PE20 tubing (inlet). The inlet is prepared from tubing with small inner diameter to minimize the dead volume (~5 μL in our case). Insert the inlet and outlet tubing into the holes drilled in the glass slide as shown in Fig. 15B.
3. Glue the tubes to the slide from the top with freshly prepared 2-part epoxy (Devcon #14250), allow the epoxy to cure for 5-10 min (Fig. 15B). This slide-tubing assembly can be prepared in advance in batches.
4. Using a sharp scalpel, trim the tubing that protrudes through the slide, so the tubing is now flush with the glass (Fig. 15C).
5. Prepare a mask from double-sided tape (Grace Bio-Labs #SA-S-1L). Although masks can be cut using a sharp scalpel, we use a laser cutter for convenience and consistency (Flux Beamo 30W). Each mask is a 24 mm by 20 mm rectangle with a 2 mm x 10 mm channel cutout in the center (Fig. 15C). These masks can be prepared in advance and stored for up to 1 year.
6. Attach the double-sided tape mask to the flush side of the glass slide assembly, aligning the channel in the mask with the holes in the slide (Fig. 15C). With the protective cover still attached to one side of the tape, use flat-head plastic tweezers (Fisher Scientific #50-238-03) to massage out all air bubbles trapped between the tape and the slide.
7. Use a diamond scribing pen (Ted Pella #54468) to cut a PEGylated coverslip into 3 equally segments, each 24 mm by 20 mm. Remove the top protective cover from the double-sided tape and attach the coverslip segment with the PEGylated side facing the double-sided tape (Fig. 15D). Use plastic tweezers to force out any air pockets trapped between the tape and the coverslip.
8. Seal the perimeter of the coverslip with freshly prepared 2-part epoxy (Fig. 15E). Allow the epoxy to cure for 5-10 min at room temperature.
9. Mount the fully assembled flow cell onto the microscope stage and, optionally, use tape to immobilize it to the stage adapter (in addition to the standard metal clips) (Fig. 15F). Connect the outlet to the syringe pump via a metal connector prepared by trimming a 21-gauge hypodermic needle (BD #305167).

**Figure 15.**
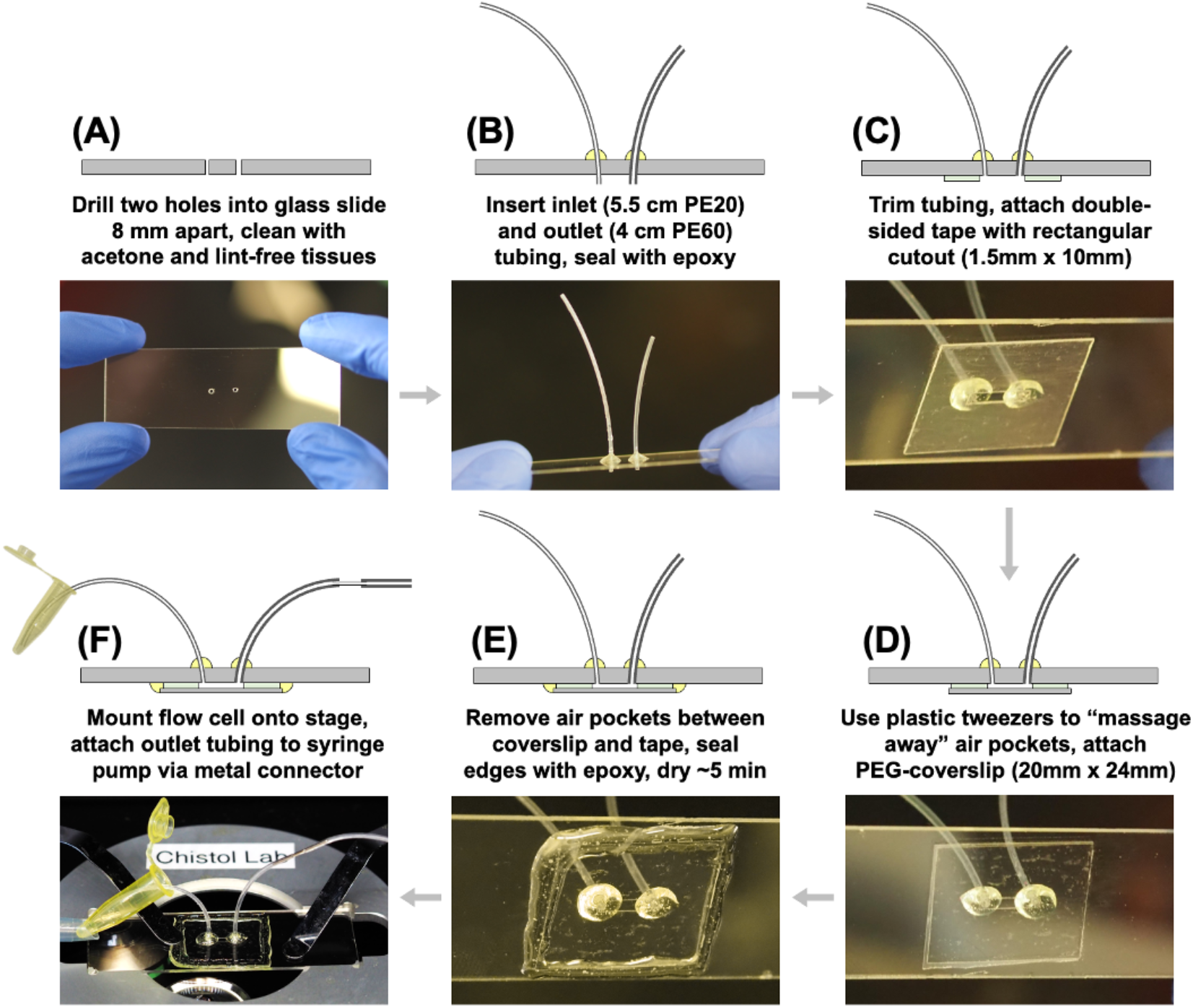
Assembling a Microfluidic Flow Cell for KEHRMIT Experiments. **(A)** Two holes are drilled into a glass slide. **(B)** Inlet and outlet tubing is inserted into each hole and secured using epoxy. **(C)** The protruding excess tubing is trimmed using a sharp scalpel (not shown) and a double-sided tape mask is attached to the bottom of the slide. **(D)** A PEGylated glass coverslip is attached to the bottom with the PEG side facing up. **(E)** The coverslip edges are sealed using epoxy. **(F)** The fully assembled flow cell is secured onto the microscope stage and connected to the syringe pump.

### Configuring the Microscope for a KERHMIT Experiment

Before beginning immunodepletions, turn on the microscope, lasers, and temperature-control enclosure set to 24 °C. It takes ~1 hour for instrument to thermally equilibrate. Open the Nikon Elements control software to begin cooling the EMCCD to −70 °C and verify the optical configurations for each color channel. Verify the dichroic and emission filters installed in the TwinCam image splitter (if using the dual-camera configuration). Before mounting the flow cell, image the reference bead slide and mechanically align the color channels using the X-shift and Y-shift micrometers on the TwinCam (only if using the dual-camera configuration). Take reference images of the bead slide in all the relevant color channels – this can be used later to align the color channels with sub-pixel precision. Use the bead slide to adjust the field aperture which clips the excitation beam to only cover the field of view – this prevents unnecessary light exposure (and photobleaching) of areas that are not being imaged.

On our microscope, the TIRF angle is set such that the TIRF controller setting of 0 corresponds to the laser light exiting the objective vertically. The laser light undergoes total internal reflection when the TIRF controller is set to ~8400 or higher (these are arbitrary units, 8400 corresponds to a beam incidence angle of ~61°). TIRF illumination can be achieved when the TIRF angle controller is set between ~8400 and ~9100, with a higher number corresponding to a shallower evanescent field. The bead slide also provides “landmarks” for fine-tuning the motorized TIRF angle illuminator. For example, when using low laser power (0.01-0.05 mW out of the objective), 0.1 μm TetraSpeck beads (Thermo Fisher #T7279) appear the brightest at a TIRF controller setting of ~8700. Double-tethered DNA stretched to ~90% of their contour length and stained with SYTOX Green in DNA buffer appear the brightest at a TIRF controller setting of ~8600. To maximize the signal-to-noise ratio, we carry out KEHRMIT experiments with the TIRF controller set to 8700-8800.

### Double-Tethering DNA Substrates in the Flow Cell

We mount the flow cell onto the microscope stage and incubate the flow cell with streptavidin during the 2^nd^ depletion round. We begin tethering the DNA immediately after starting the 3^rd^ round of depletions. All buffers are first degassed under vacuum for at least 30 min. The streptavidin solution can be optionally supplemented with fluorescently labeled streptavidin-Alexa Fluor 647 (Thermo Fisher #S21374) as a fiducial marker for focusing and computational drift correction after acquision.

Previously, we and others have supplemented the DNA solution with chloroquine to achieve greater end-to-end stretching of the DNA during tethering (Sparks and Chistol et al., 2019; Yardimci, Loveland, et al., 2012). However, we observed that chloroquine sometimes causes DNA to become cross-linked to the surface of the coverslip, so we phased out its use. To facilitate flow-stretching of long linear DNA substrates in the absence of chloroquine, we supplement the DNA buffer with 15% glycerol (see below). This, coupled with flow rates of 150 μL/min result in double tethered lambda DNAs that are stretched to 80-90% of their contour length of 16.5 μm. Finally, the SYTOX Green DNA stain (or SYTOX Orange) was selected because it is easily washed off unlike other DNA-staining dyes, as pointed out by Yardimci, Loveland, et al., 2012.

1. <u>Prepare DNA tethering solutions in 0.65 mL microcentrifuge tubes</u>
  A. Flow cell purging buffer: 525 μL DNA buffer + 25 μL 10% Tween20 detergent
  B. Streptavidin solution: 70 μL of 0.15 mg/mL streptavidin (Sigma #S4762-1mg)
  C. Flow cell purging buffer: 525 μL DNA buffer + 25 μL 10% Tween20
  D. DNA solution: 525 μL DNA buffer with 15% glycerol + 25 μL of biotinylated lambda DNA (~50ng)
  E. Wash buffer: 200 μL DNA buffer
  F. DNA stain buffer: 200 μL DNA buffer + 0.5 μL of 50 μM SYTOX Green (Thermo Fisher #S7020)
  G. DNA destain buffer: 200 μL 1xELB-sucrose
2. <u>Mount the fully assembled flow cell onto the microscope stage adapter (see Fig. 15F)</u>
  ○ Secure the flow cell with stage clips
  ○ Attach the outlet to the syringe pump
3. <u>Tether the DNA by flowing the buffers prepared earlier through the flow cell</u>
  ○ Flow mix A: 500 μL at 500 μL/min, vigorously flick the outlet tube to dislocate air bubbles
  ○ Flow mix B: 50 μL at 10 μL/min, incubate for 5-10 min after starting flow
  ○ Flow mix C: 500 μL at 500 μL/min, vigorously flick the outlet tube to dislocate air bubbles
  ○ Flow mix D: 500 μL at 150 μL/min to flow-stretch and double-tether the DNA
  ○ Flow mix E: 100 μL at 20 μL/min to wash off unbound DNA
  ○ Flow mix F: 20 μL at 20 μL/min to stain DNA
  ○ Image tethered DNA substrates and select the fields of view for KEHRMIT (see below)
  ○ Flow mix G: 150 μL at 10 μL/min to destain the DNA
4. <u>Visualize double-tethered DNA and select the fields of view</u>
  ○ Select the appropriate emission filter for SYTOX Green (Chroma #ET525/50)
  ○ Set TIRF angle (~8600 on our Nikon TIRF controller, see discussion above)
  ○ Set the excitation laser (488 nm) power to ~0.05 mW out of the objective
  ○ Set EMCCD EM Gain (between 100 and 300)
  ○ Use 100 msec exposure settings to focus on the tethered DNA, engage the focus lock system
  ○ Navigate toward the inlet in the flow cell. At the inlet the DNA density is the highest and DNA molecules diverge in all directions (see Fig. 16A).
  ○ Select the fields of view (FOVs) to be imaged during the KEHRMIT experiment. Typically, we select a grid of 6 rows by 4 columns for a KEHRMIT experiment with a 20 sec time resolution. Image the tethered DNA molecules for reference (see Fig. 16). Typically, we set the distance between FOVs to 0.1 mm (each field of view is 82 μm by 82 μm when using a 100x objective with the iXon EMCCD).

**Figure 16.**
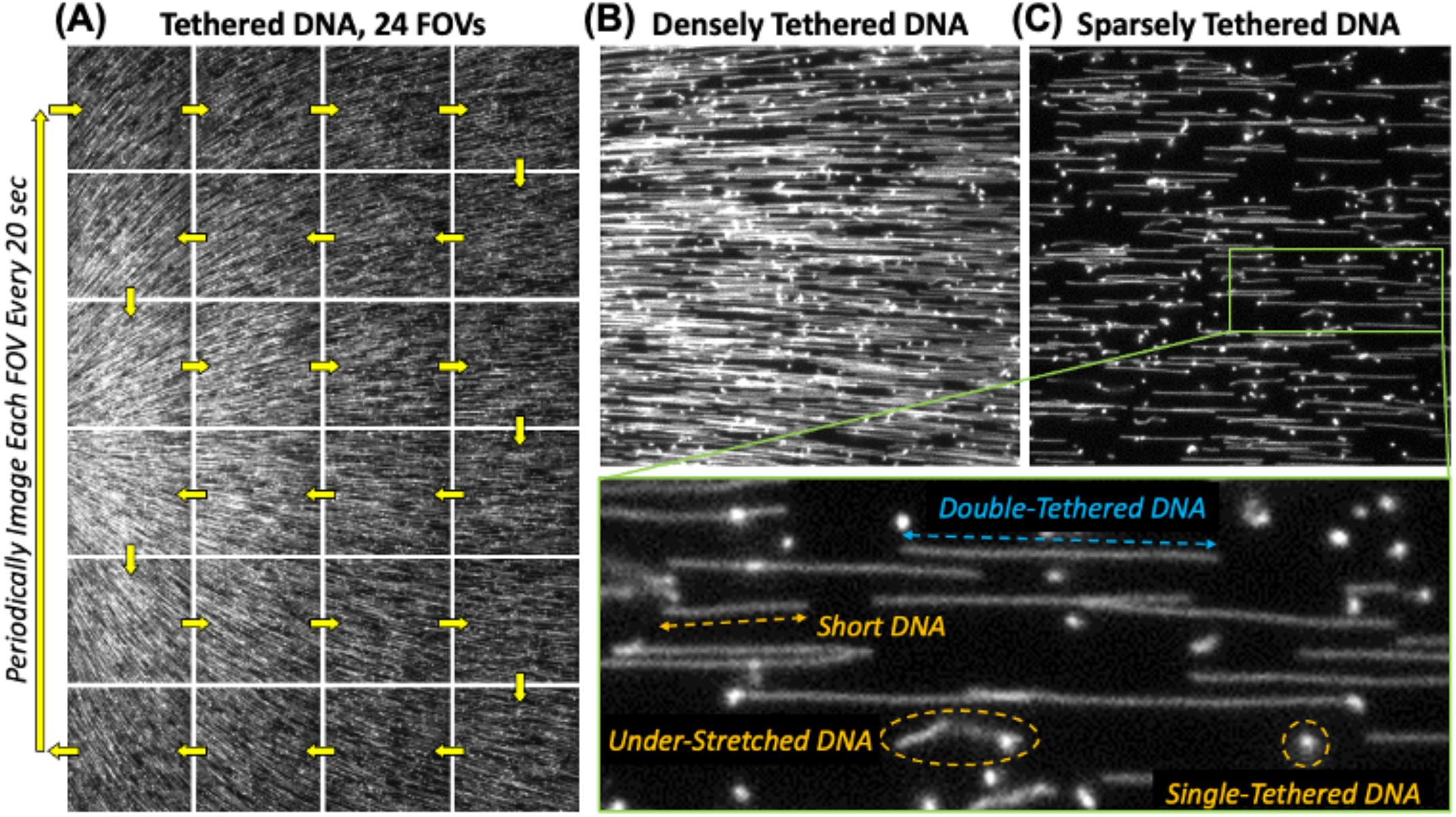
Double-Tethering Linear DNA Templates. **(A)** The DNA template with biotinylated ends is stretched via laminar flow and attaches to the streptavidin-coated surface following the fluid flow lines emerging from the flow cell inlet. During a typical KEHRMIT experiment, 24 FOVs (6 rows x 4 columns) can be periodically imaged every ~20 sec for a time course of up to 2 hrs. **(B-C)** The DNA density depends on the proximity to the inlet – allowing the researcher to choose optimal fields of view. Panels B and C illustrate the upper and lower limit of acceptable densities of tethered DNA. **(C, inset)** We routinely observe four classes of DNA molecules immobilized onto the coverslip: (i) properly stretched double tethered DNA of the correct size (blue arrows); (ii) properly stretched double tethered DNA molecules that are too short (orange arrows); (iii) under-stretched DNA molecules with a lot of slack (orange oval); and (iv) DNA molecules that are tethered to the coverslip only at one end (orange circle). Only DNA substrates from class (i) are suitable for KEHRMIT imaging.

### Performing the KEHRMIT Experiment

Typically, the replication initiation reaction contains too much fluorescent GINS (100-300 nM), so imaging is performed during the replication elongation step. In contrast to the biochemical assay where DNA is licensed for 30 min, KEHRMIT licensing is performed for only 2-5 min to precisely control the number of dormant origins. The number of origins that fire is also controlled by adjusting the concentration of GINS^AF647^ in the replication initiation mix, and the duration of the replication initiation incubation step. All the incubation times are measured from the moment the flow is turned on.

1. <u>Configure the Nikon Elements software for a high-throughput KEHRMIT experiment</u>
  ○ multi-point (24 FOVs)
  ○ multi-wavelength (488 nm and 647 nm)
  ○ multi-timepoint (1 frame every 30 sec for a total of 120 frames = 1 hour long experiment)
2. <u>Verify the optical configurations for each color channel</u>
  ○ 488 nm channel (CIDER imaging of Fen1^mKikG^)
    - EM Gain 300
    - 100 msec exposure
    - ~0.1 mW laser power (out of the objective)
    - TIRF illuminator setting of ~8800 (~64° angle of incidence)
  ○ 647 nm channel (KEHRMIT imaging of GINS^AF647^)
    - EM Gain 300
    - 100 msec exposure
    - ~0.25 mW laser power (out of the objective)
    - TIRF illuminator setting of ~8800 (~64° angle of incidence)
3. <u>Introduce extract into the flow cell to replicate DNA</u>
  I. Flow the DNA licensing mix: 15 μL at 10 μL/min, incubate for 3 min after starting flow
    ○ During licensing, focus on the streptavidin-Alexa Fluor 647 fiducials immobilized on the surface, and add a focusing offset of 0.3-0.4 μm to compensate for the fact that tethered DNA molecules are above the glass coverslip surface, as discussed above.
  II. Flow the DNA replication initiation mix: 25 μL at 10 μL/min, incubate for 3 min after starting flow
  III. Flow the DNA replication elongation mix: 55 μL at 10 μL/min
4. <u>Acquire data for 60 minutes</u>
  ○ Begin imaging 2 min after starting the flow of the DNA replication elongation mix
5. <u>Dis-assemble flow cell, clean oil from objective, purge syringe pump, turn off instrument.</u>

### Performing Data Analysis

Here we summarize the data analysis workflow for a simple two-color KEHRMIT+CIDER experiment. The goal of this pipeline is to generate kymograms for each DNA molecule that is undergoing replication. In each kymogram, we can then identify and select regions of interest (ROIs) that can further be analyzed with specific measurements in mind (determining helicase speed, detecting helicase pausing events, detecting changes in helicase speed, etc.). Figures 17 and 18 outline the data analysis workflow for KEHRMIT and CIDER/PhADE respectively. The basic analysis code and a sample data set is available on GitHub at https://github.com/ChistolLab/KEHRMIT.

1. Each single molecule experiment is performed at least two times, preferably with different extract preparations to account for extract-to-extract variability.
2. Single-molecule data is acquired using the Nikon NIS Elements software and exported as a multi-page TIFF for each field of view. These files are saved in a folder named “Exported TIFFs”.
3. Each TIFF file is subjected to drift-correction in ImageJ using the Image Stabilizer plugin (written by Kang Li, Carnegie Melon University) and our own ImageJ macro to automate the process (Fig. 17A, 18A). These files are saved in a folder named “Stabilized TIFFs”.
4. The drift-corrected TIFF files are subsequently processed in MATLAB using our own KEHRMIT package (Fig. 17-18). The default mode is 2-color analysis with a KEHRMIT channel and a CIDER channel, but the code can also be used in single-channel only mode (KEHRMIT only or CIDER only).
5. First, the drift-corrected TIFF file is opened, background subtraction is performed to remove non-uniform illumination. Next the maximum projection of each channel is generated – this illustrates the highest brightness of each pixel over the course of the movie. The maximum projection is a convenient way of visualizing the extent of replicated DNA (Fig. 17B, 18B). Individual DNA molecules are then selected manually by drawing a line over each “bright track” in the maximum projection representation (Fig. 17C, 18C). The code then automatically generates kymograms for each molecule selection and saves the results of this intermediate analysis as a MAT file. These tasks are accomplished by running the “SelectMolecules” function.
6. Second, the kymograms generated in the previous step (Fig. 17D, 18D) are inspected to identify regions of interest (ROIs) using the “SelectKymoROI” function. ROIs may be regions where a replication origin fires bidirectionally, or where two replisomes converge and undergo replication termination, or where a replisome encounters a DNA lesion and pauses. The spatial and temporal extent of the ROI is then defined by the user and saved by the program for downstream analysis. More than one type of ROI can be defined by the user for custom analysis workflows.
7. Downstream analysis of the ROI selections may include the following:
  ○ Measuring the extend of the replicated DNA in the CIDER channel to compute replication fork speed
  ○ Fitting the CMG helicase signal to a point-spread function to track helicase movement
  ○ Measuring helicase pausing data, or the lifetime of specific intermediate states (pausing)

**Figure 17.**
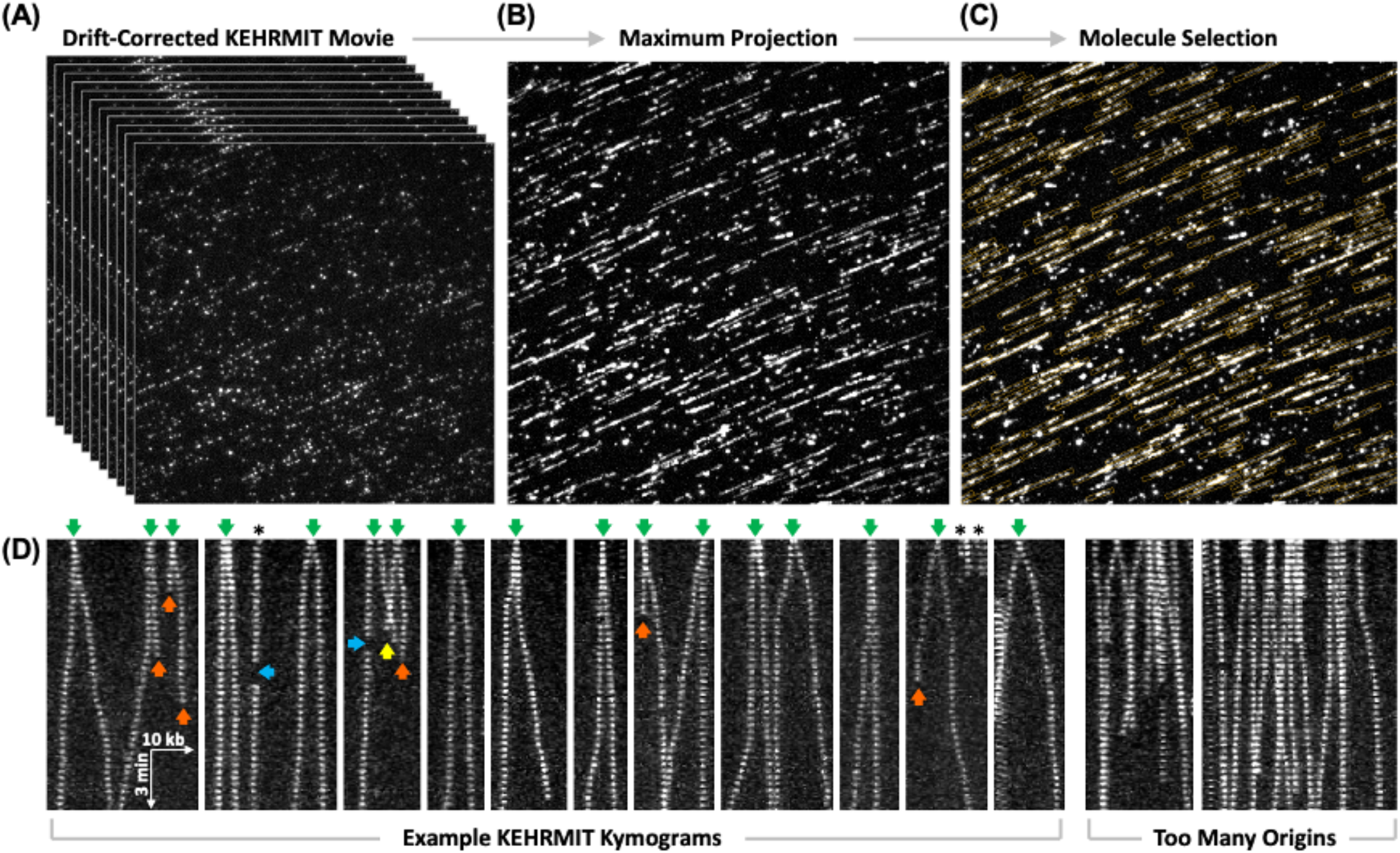
KEHRMIT Analysis Workflow. KEHRMIT movies are drift-corrected **(A)**, and a maximum projection is calculated **(B)**. Contiguous trajectories in the maximum projection represent tracks of processive helicase movement. **(C)** These tracks are selected for further analysis. Kymograms corresponding to each selection are automatically generated. **(D)** Example KEHRMIT kymograms illustrating bidirectional DNA replication from individual origins (green arrows). Orange arrows point to events where the fluorophore photobleached. Blue arrows indicate blinking events, where the fluorophore temporarily switches to the dark state and back. The yellow arrow points to a replisome convergence event and subsequent unloading of the replicative helicases involved. If the density of replication origins is too high, it becomes challenging to analyze and interpret the data.

**Figure 18.**
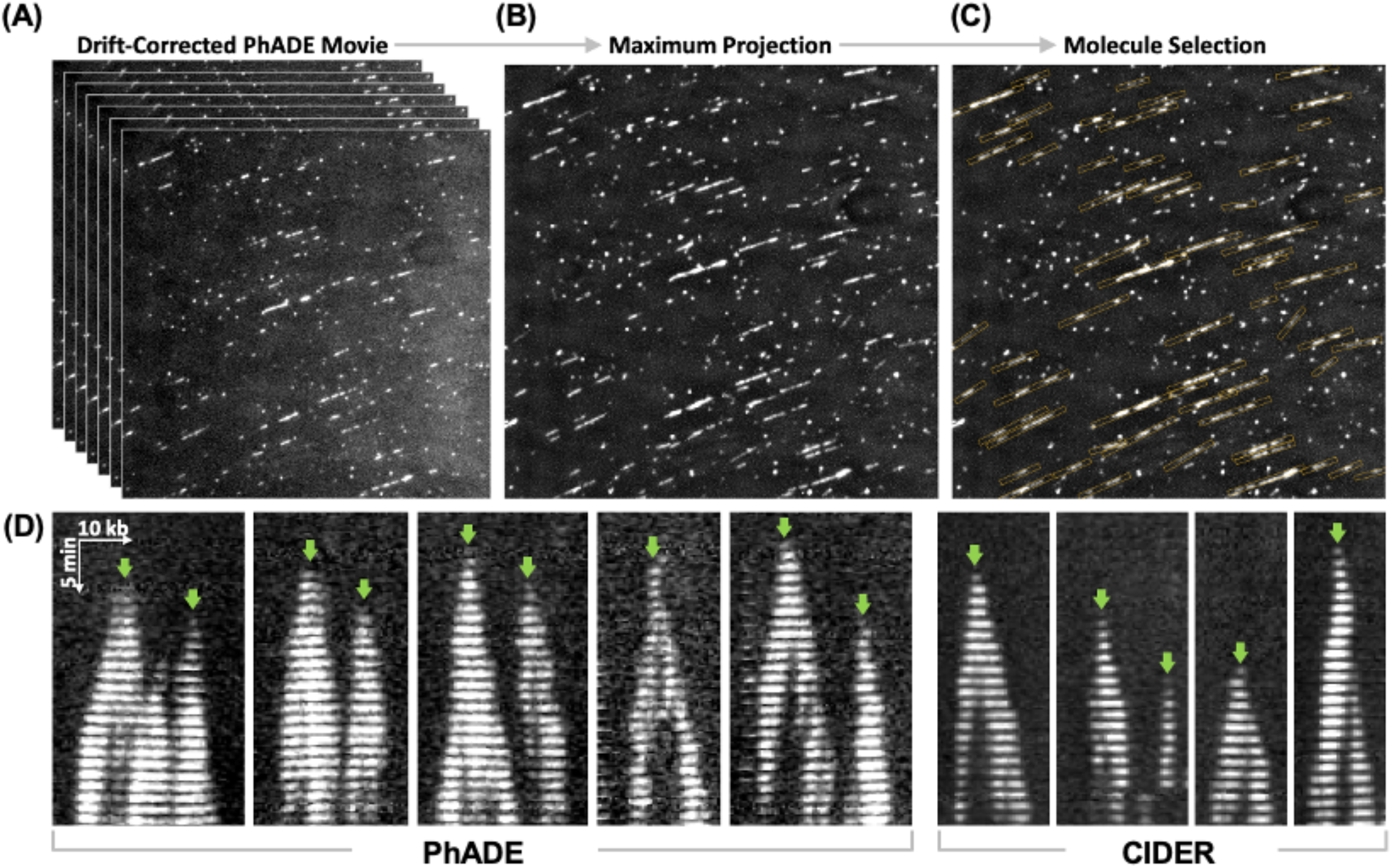
PhADE or CIDER Analysis Workflow. PhADE and CIDER movies are analyzed using the exact same workflow. **(A)** The movie is drift-corrected, and a maximum projection is calculated **(B)**. Contiguous trajectories in the maximum projection represent tracks of processive DNA replication. **(C)** These tracks are selected for further analysis and kymograms are automatically generated for each selection. **(D)** Example kymograms from PhADE and CIDER experiments illustrates a comparable signal-to-noise performance. Green arrows indicate replication initiation events. Although PhADE and CIDER appear to have a similar SNR in these kymograms, each PhADE exposure was 100 msec whereat each CIDER exposure was 500 msec. Both PhADE and CIDER data was collected using Fen1^mKikGR^. For PhADE, the 405 nm and 561 nm lasers were used for photoactivation and photoexcitation respectively. For CIDER, the 488 nm laser was used for direct imaging.

### Conclusions

In recent years, single-molecule imaging of eukaryotic DNA replication has provided invaluable insights into this fundamental biological process. In particular, KEHRMIT is an attractive approach as it leverages nuclear egg extracts to recapitulate metazoan DNA replication without the need for a complete biochemical reconstitution. The goal of this article is to facilitate the adoption of our approach by other laboratories and to inspire new single-molecule studies in the field. To this end we provide detailed protocols and illustrations of key steps in our workflow as well as technical context that is beyond the scope of a traditional research article. Please do not hesitate to contact us for additional technical details.

## Acknowledgements

We thank members of the Chistol lab (Luke Lynch, Riki Terui, Dhruva Deshpande, Larissa Sambel, and Jinho Park) for suggestions and feedback. Scott Berger is supported by the Stanford Graduate Fellowship (SGF) and the New Science fellowship. Gheorghe Chistol is supported by an NSF CAREER Award (2144481) and an NIGMS R35 award (GM147060).

## Author Contributions

S.B. and G.C. wrote the manuscript and compiled the figures. G.C. wrote the data analysis code.

